# Depth-dependent dynamics of protist communities as an integral part of spring succession in a freshwater reservoir

**DOI:** 10.1101/2024.02.01.578394

**Authors:** Indranil Mukherjee, Vesna Grujčić, Michaela M Salcher, Petr Znachor, Jaromír Seďa, Miloslav Devetter, Pavel Rychtecký, Karel Šimek, Tanja Shabarova

## Abstract

**Background:** Protists are essential contributors to eukaryotic diversity and exert profound influence on carbon fluxes and energy transfer in freshwaters. Despite their significance, there is a notable gap in research on protistan dynamics, particularly in the deeper strata of temperate lakes.

This study aimed to address this gap by integrating protists into the well-described spring dynamics of Římov reservoir, Czech Republic. Over a two-month period covering transition from mixing to established stratification, we collected water samples from three reservoir depths (0.5, 10 and 30 m) with a frequency of up to three times per week. Microbial eukaryotic and prokaryotic communities were analysed using SSU rRNA gene amplicon sequencing and dominant protistan groups were enumerated by Catalysed Reporter Deposition-Fluorescence *in situ* Hybridization (CARD-FISH). Additionally, we collected samples for water chemistry, phyto- and zooplankton composition analyses.

**Results:** Following the rapid changes in environmental and biotic parameters during spring, protistan and bacterial communities displayed swift transition from a homogeneous community to distinct strata-specific communities. Epilimnion exhibited the prevalence of auto-, mixotrophic protists dominated by cryptophytes and associated with spring algal bloom-specialized bacteria. In contrast, meta- and hypolimnion showcased the development of protist community dominated by putative parasitic Perkinsozoa, detritus or particle-associated ciliates, cercozoans and excavate protists co-occurring with bacteria associated with lake snow.

**Conclusions:** Our high-resolution sampling matching the typical dividing time of microbes along with the combined microscopic and molecular approach and inclusion of all the components of microbial food web allowed us to follow depth-specific populations’ successions and interactions in a deep lentic ecosystem.

## Background

Microbial eukaryotes, the main contributors of eukaryotic diversity on Earth, include a remarkable diversity of single-celled planktonic protists that are omnipresent across all aquatic environments. Their significance in ecosystems stems from their diverse functions in carbon fluxes and energy transfer in aquatic food webs[1, 2], encompassing a spectrum of ecological and biochemical roles and nutrition modes, from autotrophy/mixotrophy to heterotrophy (including predation, decomposition, parasitism, and osmotrophy)[3–7].

For a long time, research on protists faced limitations due to labour-intensive microscopic analyses and the often low morphological resolution of these organisms, particularly evident in the case of numerically dominant but taxonomically diverse small cells of heterotrophic nanoflagellates (HNF)[8–10]. However, the situation changed with the advent of high-throughput sequencing and the accessibility of 18S rRNA gene amplicon approaches. This transformation propelled protist research into a prominent focus within microbial ecology. Subsequently, there has been a rapid increase in studies from the early 2000s reporting an unprecedented diversity of nano-sized protists in marine[11–15] and freshwater[16–22] environments.

Most studies conducted in freshwaters have focused on the epilimnion, the upper water layer of highest productivity, while protist communities of deeper strata have been largely neglected, with only a few exceptions[21, 23, 24]. Additionally, the temporal resolution of most datasets is limited to low sampling frequencies (weeks to months) that is insufficient for capturing the rapid dynamics of fast-growing protists (replicating in hours to days)[7, 25, 26]. This considerably hinders the understanding of dynamic environmental events such as phytoplankton spring blooms[27, 28] or impacts of dramatic environmental disturbances[29]. In temperate freshwater lakes, the onset of spring is marked by a physical mixing event after ice melt, which uniformly distributes microbial populations in the water column. The increase of light intensity and air temperature leads to a thermal stratification of the water column and rapid growth of phototrophic organisms in the epilimnion that serve as a base for the aquatic food web. The succession of phytoplankton and zooplankton during this phase was well described in the original and revised Plankton Ecology Group (PEG) model[30, 31].

Recently, an attempt was made to expand the PEG model to encompass seasonal dynamics of prokaryotes[32]. The detailed resolution of bacterial spring dynamics showed a dominance of fast-growing phytoplankton bloom-associated Bacteroidota and Gammaproteobacteria at the beginning of spring bloom[27, 28, 33, 34], succeeded by small-sized Actinobacteriota resistant to protistan grazing[27, 35]. Even though protistan abundance was included in the revised PEG model[31] and protists were recorded in various prokaryotic studies for the estimation of top down control, there is a lack of information on the dynamics of individual protistan populations, their ecological associations and functions within the aquatic food web in distinct lake strata[36].

Metagenomic analyses which successfully resolve bacterial populations, have an unfortunate constraint for research on protist communities as computational power and necessary depth of sequencing grow exponentially with the size of sequenced organisms[28]. Therefore, many studies introduce arbitrary cutoffs of 20 or 5 µm for filtration and thus do not provide information on larger protists as well as symbiotic interactions, which might be crucial during the springtime[30, 31].

In this study conducted in the temperate freshwater Římov reservoir (Czech Republic), we used a cutoff of 200 µm for biomass collection and hybrid approach combining 18S rRNA gene amplicon sequencing with the fluorescent labelling technique Catalysed Reporter Deposition-Fluorescence *in situ* Hybridization (CARD–FISH)[6, 22, 24, 37, 38]. This allowed us to visualise and enumerate specific protistan lineages, which dominated the sequencing data[36, 39]. Additionally, we analyzed 16S rRNA gene amplicons of the prokaryotic community, phyto- and zooplankton, viruses, and chemical parameters, aiming at identifying dominant protist populations and their major interactions to refine our comprehension of the spring plankton succession in different strata of the reservoir.

## Methods

### Study site and sampling procedure

The Římov reservoir, situated in South Bohemia, Czech Republic, is a dimictic, meso-eutrophic canyon-shaped reservoir which serves as an important drinking water supply. It covers an area of 2.06 km^2^ with a volume of 34.5×10^6^ m^3^ and an average summer retention time of 77 days. It has been studied since 1979 at well–established stations [40], one of them located in the lacustrine zone near the dam (48.8475817N, 14.4902242E, max. depth 42 m) was used for sampling in our study.

We conducted high-frequency sampling, up to three times per week during the period between 31^st^ March to 25^th^ May 2016. We sampled 3 depths (0.5, 10 and 30 m), which corresponded to the epi-, meta-, and hypolimnion, respectively. The samples were taken with a Friedinger sampler (Šramhauser; spol.s.r.o., Dolní Bukovsko, Czech Republic). For each depth ten litres of water were prefiltered through a 200 µm mesh plankton net into a plastic barrel which was cleaned with household bleach and rinsed with Milli-Q and sample water. Physical and chemical parameters, i.e., water temperature, pH, dissolved oxygen, and oxygen saturation were measured with a multiparametric probe YSI EXO2 (Yellow Springs Instruments, Yellow Springs, OH, USA). A submersible fluorescence probe (FluoroProbe; bbe–Moldaence, Kiel, Germany) was employed to measure chlorophyll *a* (Chl-*a*) concentrations at 0.2 m intervals down till 20 m depth. Water transparency was measured using Secchi disc. Samples for chemical analysis were collected in separate bottles.

Phytoplankton samples were collected from 0.5 m depth and preserved with a Lugol’s solution for further processing. Crustacean zooplankton was sampled once a week by vertical hauls using an Apstein plankton net (200-um mesh). Net hauling provided an integrated sample for the upper 5 m water column representing the epilimnetic layer. Two hauls were combined into one sample and preserved with formaldehyde (4% final concentration) for subsequent processing in the laboratory. Small sized rotifers were sampled analogically once a week as integrated sample from the upper 5 m water column using a plastic tube of the appropriate length. Subsequently, a volume of 40 l of collected water was quantitatively filtered using a 35 µm thickening net. The collected material was preserved with formaldehyde (4% final concentration).

### Chemical analysis

Samples were analysed for pH, dissolved organic carbon (DOC), dissolved nitrogen (DN), dissolved silica (DSi), total phosphorus (TP), dissolved phosphorus (DP), dissolved reactive phosphorus (DRP), NH_4_-N, NO3-N and absorbance was measured at wavelengths 254, 300, 350 and 400 nm (**Additional file 1)** using methods summarized in Znachor et al[40].

### Enumeration of microbial cells, viruses, phytoplankton and zooplankton

For each depth, subsamples were fixed with the Lugol-formaldehyde-thiosulfate decolourization technique (2% final concentration of formaldehyde) to minimize ejection of protistan food vacuole contents[41]. These samples were used for the enumeration of bacteria on black 0.2 μm pore-size filters (Osmonics, Inc., Livermore, CA, USA) and eukaryotes (flagellates and ciliates) on black 1 μm pore-size filters. All samples were stained with DAPI (4′, 6-diamidino-2-phenylindole, 1 μg ml^-1^ final concentration) and microbes were counted via epifluorescence microscopy (Olympus BX 53; Optical Co., Tokyo, Japan). For enumeration of virus-like particles (VLP), subsamples were fixed with glutaraldehyde (1% final concentration) for 10 min and flush-frozen with liquid nitrogen and stored at -80°C until further processing. VLPs were counted with an inFlux V-GS cell sorter (Becton Dickinson, Franklin Lakes, NJ. USA) as previously described[42].

Phytoplankton species were enumerated employing the Utermöhl method under an inverted microscope (Olympus IX 71)[43]. The mean dimension of algal cells were obtained for biovolume calculation using the approximation of cell morphology to regular geometric shapes[44].

For the analysis of zooplankton, formaldehyde from the preserved material was removed and partially replaced by tap water. Further processing was performed by a classical microscopical counting of different species with a series of species determination keys[45]. Rotifer abundance was analysed in exact subsamples in counting chamber using dissecting microscope Leica DM 2500 under magnification of 25-40×. Species determination was done according to Koste 1978[46] in light of more recent literature in particular families and recent taxonomy.

### DNA extraction and sequencing

Prokaryotic and eukaryotic biomass was collected on 0.2 µm pore-size filters (47 mm diameter; Osmonics, Minnetonka, MN, USA) from 1500-2000 ml of water. DNA was extracted with the Power Water DNA isolation kit (MO BIO Laboratories, Inc, Carlsbad, CA, USA). Prokaryotic 16S rRNA fragments (V4 region) were amplified using primer pair 515F and 926R[47] and sequenced on an Illumina MiSeq platform (PE300) with V3 chemistry at Genome Research Core of the University of Illinois (Chicago, USA). Eukaryotic amplicons (V9 region) were prepared using Euk_1391F and EukBr–7R primers (https://earthmicrobiome.org/protocols-and-standards/18s/) and sequenced on an Illumina MiSeq platform (PE250) with V2 chemistry at SEQme company (Dobříš, Czech Republic).

### Sequence Analysis

Primers were cut from the demultiplexed reads using Cutadapt software v2.8[48]. Trimmed sequences were processed using DADA2 pipeline v1.16.0[49] with standard parameters (https://benjjneb.github.io/dada2/tutorial.html) in R (R Core Team 2020). For taxonomic identification at ASV (Amplicon Sequence Variant) level, SILVA v138[50, 51] and PR^2^ v4.14.0[52] were used for prokaryotes and eukaryotes, respectively. All irrelevant reads (mitochondria and plastids for prokaryotes and metazoa and fungi for eukaryotes) and singletons were excluded, and both datasets were rarefied to the smallest read number prior diversity estimation and statistical analysis. In order to access relatedness of microbial communities of different lake strata and their temporal dynamics, a Bray-Curtis dissimilarity distance matrices were calculated for protists and prokaryotes. Based on the obtained matrices, we performed nonparametric multidimensional scaling (nMDS) analysis using XLSTAT14 (Addinsoft, USA). Diversity estimators and indexes were calculated using vegan package in R[53]. A co-occurrence network was used to find associations between common protistan and prokaryotic ASVs (relative abundances >0.5% in at least one sample).

Subsequently, all possible pairwise Spearman’s rank correlations were calculated with the R script https://github.com/RichieJu520/Co-occurrence_Network_Analysis[54]. Only robust (|r|>0.7) and statistically significant (p<0.05) correlations were visualized in Gephi v0.9.2[55] with subsequent modular analysis. The sequence data generated from amplicon sequencing were submitted to the European Nucleotide Archive (ENA) and are available under the BioProject: PRJEB66298.

### Phylogenetic tree reconstruction, design of novel eukaryotic probes, and catalysed reporter deposition fluorescence *in situ* hybridization (CARD**–**FISH)

Representative amplicons of the 30 most abundant protistan ASVs were aligned with the SINA aligner[56] and imported into ARB[57] using the SILVA database SSURef_NR99_123[58]. Alignments were manually refined and a maximum likelihood tree (1000 bootstraps) including their closest relatives was constructed on a dedicated web server[59] (**Additional files 2–4**). Oligonucleotide probes targeting a small, monophyletic lineage of katablepharids (Kat2-651), Telonema (Telo-1250) and Novel Clade 10 of Cercozoa (NC10–1290) were designed in ARB using the tools probe_design and probe_check and evaluated with the web tool math-FISH[60]. The formamide percentages were optimized in environmental samples **(Table 1, Additional file 5)**.

**Table 1:**
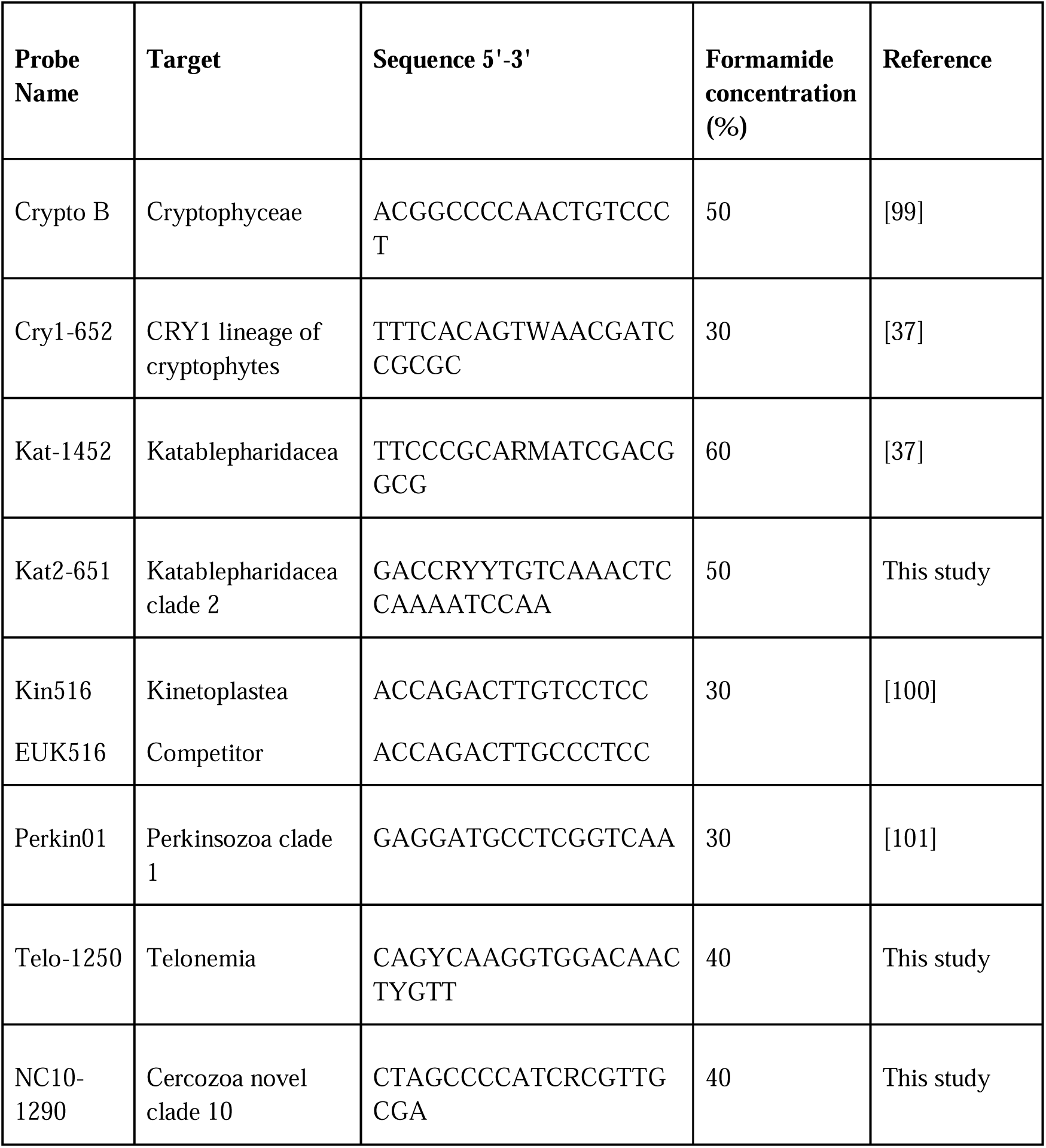
CARD-FISH probes used in the study.

CARD-FISH was carried out for eight protistan lineages (**Table 1**) following published protocols[36]. Hybridized cells were visualized with an Olympus BX 53 epifluorescence microscope under 1000× magnification at blue/UV excitations.

### Estimation of bacterivory rates of HNF and ciliates

Flagellate and ciliate bacterivory rates in the epilimnion were estimated using fluorescently labelled bacteria (FLB)[61], prepared from a mixture of strains from the genus *Limnohabitans* and *Polynucleobacter*[62]. Briefly, the FLB tracers were added to constitute 8–20% of total bacteria. Samples were incubated at *in situ* temperature with FLB tracers for 5 and 30 mins for ciliate and flagellate grazing rates, respectively. Incubation was terminated with fixation and DAPI stained subsamples were prepared as described above for microscopical analysis[62, 63]. A minimum of 100 ciliates and 200 HNF were inspected for FLB ingestion in each sample. To estimate total protistan grazing, average bacterial uptake rates of ciliates and HNF were multiplied by their *in situ* abundances.

## Results

### Physical and chemical parameters and abundance of microbes

The temperature profile from the first sampling date (31^st^ March 2016) indicated an almost homogeneously mixed water column with only two degrees difference between the surface and the bottom **(Figure 1a**). Similarly, the concentration of dissolved oxygen was almost uniform between 10 to 12 mg l^-1^ in the entire water column (**Figure 1b**). At the end of sampling campaign (25^th^ May), when the water column was stratified, the epilimnion temperature reached 16.6 °C and the thermocline was established between 5-10 m depth. Soon after the beginning of the sampling campaign, hypoxia started to develop in the bottom layers, which became anoxic at the end, however, the sampling depth of hypolimnion (30 m) was always oxygenated (>6.9 mg l^-1^; **Figure 1b**). A Chl-*a* maximum (19.6 μg l^-1^) was observed in the epilimnion in the first week of sampling after which the concentration dropped and remained low till the end of the study (**Figure 1c**).

**Figure 1:**
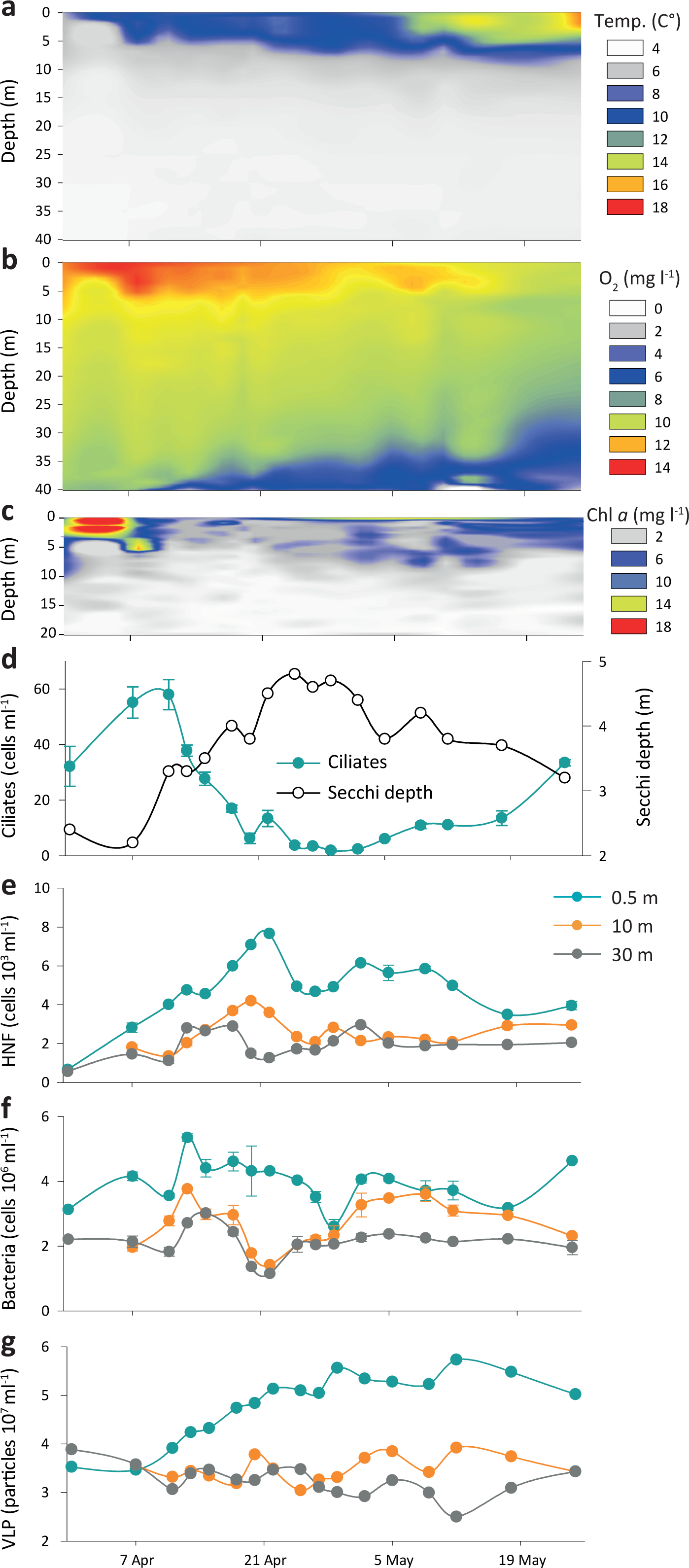
Main physical and chemical parameters and abundances of microorganisms observed in the Římov reservoir during the study. a – thermal structure of the water column (Temp). b – vertical distribution of oxygen (O_2_). c – chlorophyll *a* profile (0-20 m). d – abundances of ciliates in the epilimnion and Secchi depths. e –abundances of heterotrophic nanoflagellates (HNF) at three depths. f – bacterial abundances at three depths. g – abundances of virus like particles (VLP) at three depths.

Ciliates increased in the first two weeks to maximum abundance of 60 ind. ml^-1^ (**Figure 1d**), and were dominated by prostomes i.e., *Urotricha* spp. and *Balanion planktonicum,* classical hunters of small algae and flagellates[64]. Ciliate abundances dropped sharply during the clear water phase and recovered simultaneously with Chl-*a* concentrations at the study end. Low counts of HNF, prokaryotes and virus-like particles (VLP), were recorded at the start of the sampling (**Figure 1e, f, g**). During the first two weeks, the numbers of HNF increased in all water strata, with a maximum of 2.5×10^3^ cells ml^-1^ in the epilimnion and lower peaks in deeper layers (1.8×10^3^ and 1.3×10^3^ cells ml^-1^ at 10 m and 30 m, respectively; **Figure 1e**). Thereafter, the abundances gradually decreased in all layers with some occasional peaks. Bacterial abundances reached maxima in the epilimnion in mid-April (5.3×10^6^ cells ml^-1^), whereas numbers remained low in the metalimnion and hypolimnion (averaging 2.6×10^6^ cells ml^-1^ and 2.0×10^6^ cells ml^-1^, respectively) (**Figure 1f**). VLP steadily increased at 0.5 m to 5.6×10^7^ VLP ml^-1^ and plateaued towards the end, while abundances at 10 and 30 m remained relatively stable with maxima of 3.8×10^7^ VLP ml^-1^ at 10 m and 3.9×10^7^ VLP ml^-1^ at 30 m (**Figure 1g**).

Phytoplankton biovolume in the epilimnion peaked with 3.6 mm^-3^ l^-1^ in the first week of April, showed a sharp decline within the following two weeks and remained low until the end (**Figure 2a**). Cryptophytes (*Cryptomonas reflexa, Rhodomonas minuta*) dominated the phytoplankton throughout the sampling period together with diatoms (*Cyclotella* sp., *Fragilaria sp.*) and chrysophytes (*Chrysococcus* sp.). Rotifers and cladocerans followed the phytoplankton dynamics and showed maxima in the second week of sampling (**Figure 2b**).

**Figure 2:**
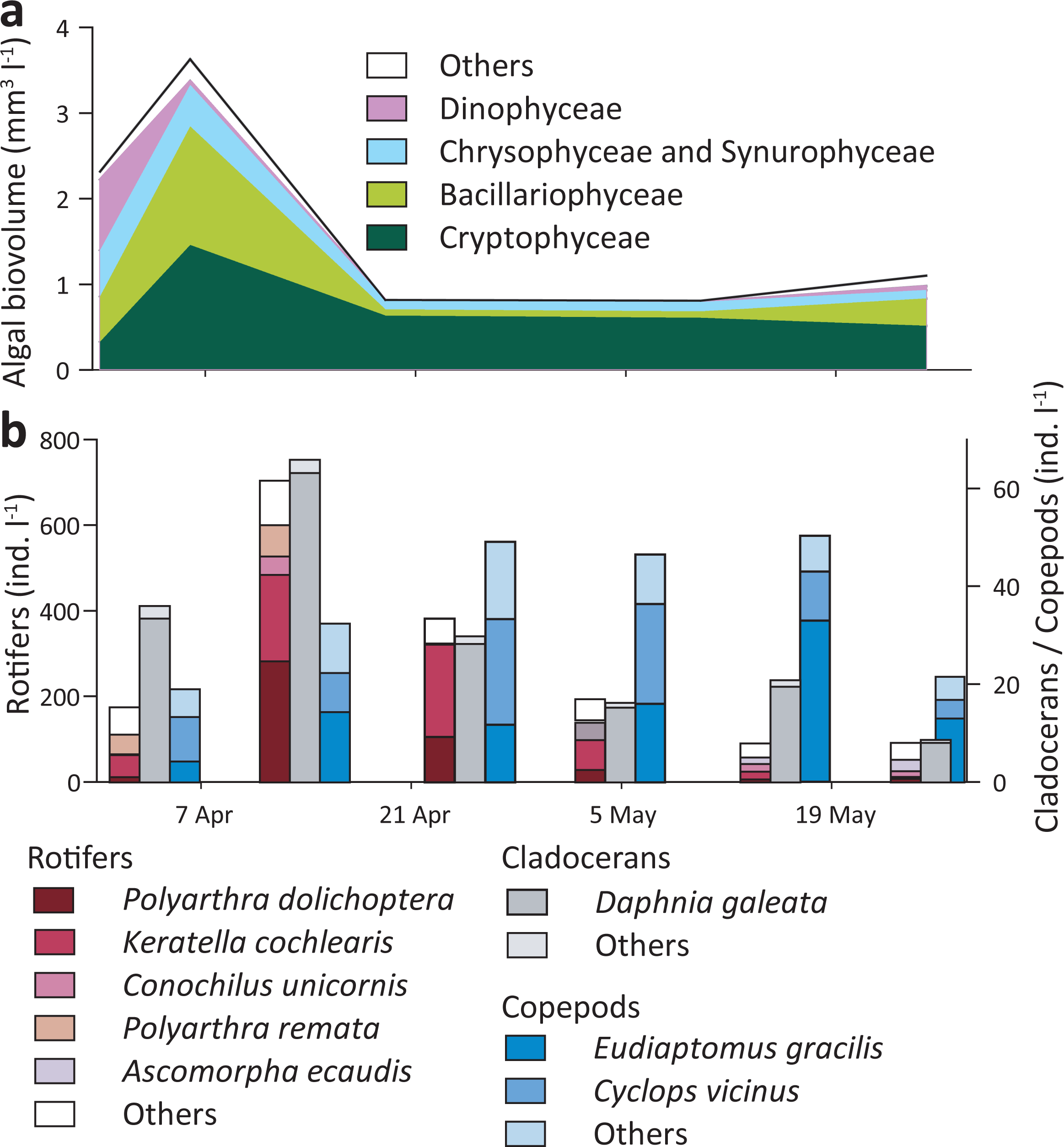
Phytoplankton biovolume (a) and zooplankton abundance (b) in the epilimnion of Římov reservoir during the study.

Copepod numbers increased slowly, but their populations remained stable in the second part of the study when densities of rotifers and cladocerans decreased.

### Community composition of microbes in the water column

From 51 DNA samples, 47 and 43 were successfully amplified and sequenced for prokaryotic and eukaryotic analyses, respectively. Datasets were rarefied to 31011 (prokaryotes) and 13202 (protists) reads per sample. Both taxonomic entities displayed similar temporal developments based on their ASV dynamics. After mixing, we observed a fast separation of epilimnion samples from the deeper lake strata and undirected fluctuations and delayed differentiation of meta- and hypolimnetic communities (**Figure 3**). Reduction of diversity over the time was noticed in both prokaryotes and protists especially in the epilimnion (**Figure 3**). Despite similar dynamic patterns observed at highly resolved taxonomic level, prokaryotes showed more unified composition at family to phylum levels than protists (**Figure 4a**). Specifically, Actinobacteriota, Verrucomicrobiota, Chloroflexota and several families of Gammaproteobacteria (Comamonadaceae, Burkholderiaceae and Methylophilaceae) were distributed at comparable relative abundances in all samples from all layers. In contrast, among protists only Katablepharida and Ciliophora showed comparable distributions. (**Figure 4b, Additional file 6**).

**Figure 3:**
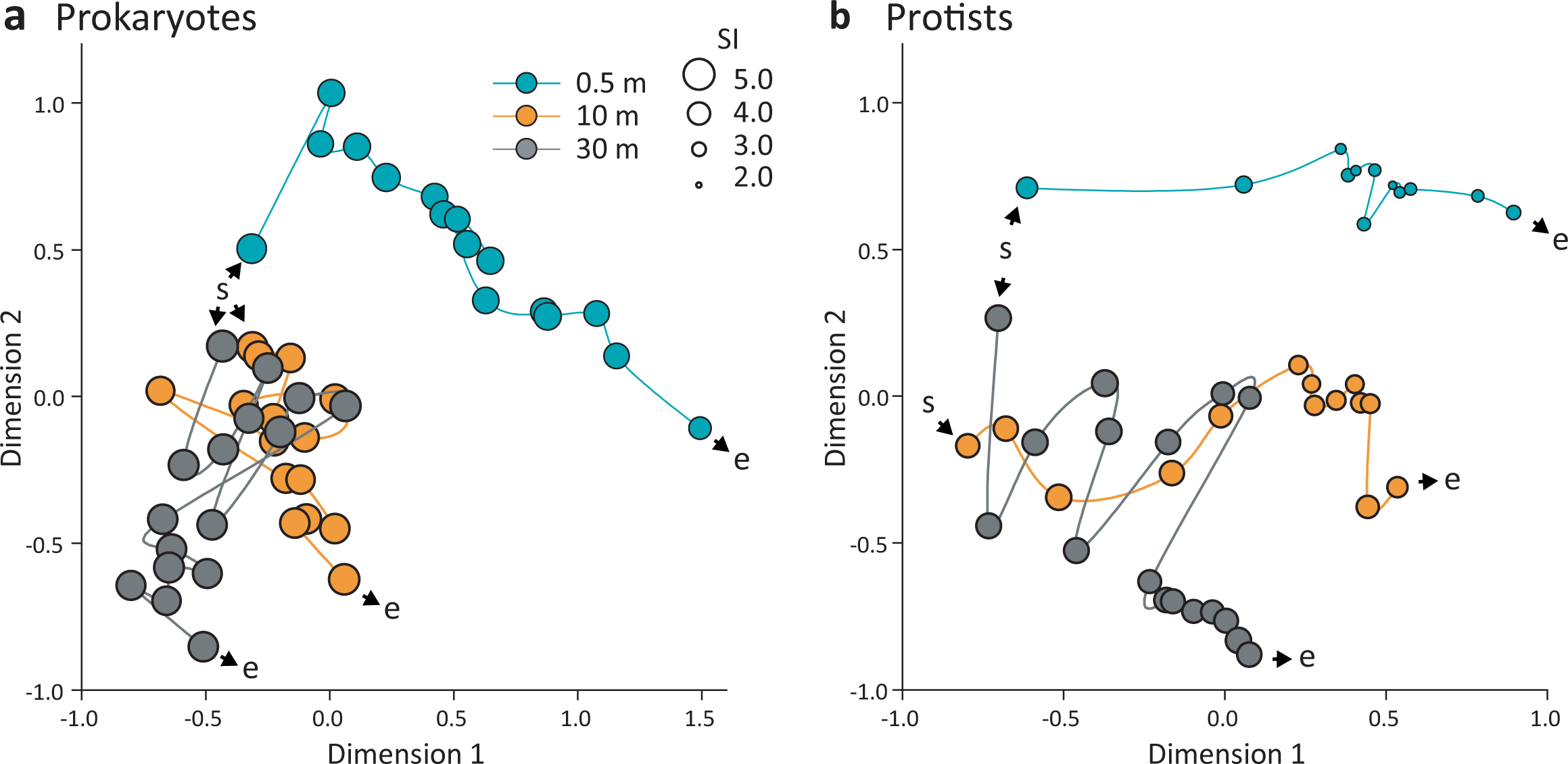
Nonparametric multidimensional scaling plots reflecting dynamics of prokaryotic (a) and protistan (b) communities at three depths of Římov reservoir. Plots are based on Bray-Curtis’s distance matrices calculated on rarefied ASV datasets. Kruskal’s stress values are 0.096 and 0.079 for prokaryotic and protistan plots respectively. The sampling start is indicated with the letters ‘s’, and the end is indicated with the letters ‘e’. The lines are connecting communities according to the temporal course of the study. Diameters of the circles correspond to the Shannon-Wiener diversity index (SI).

**Figure 4:**
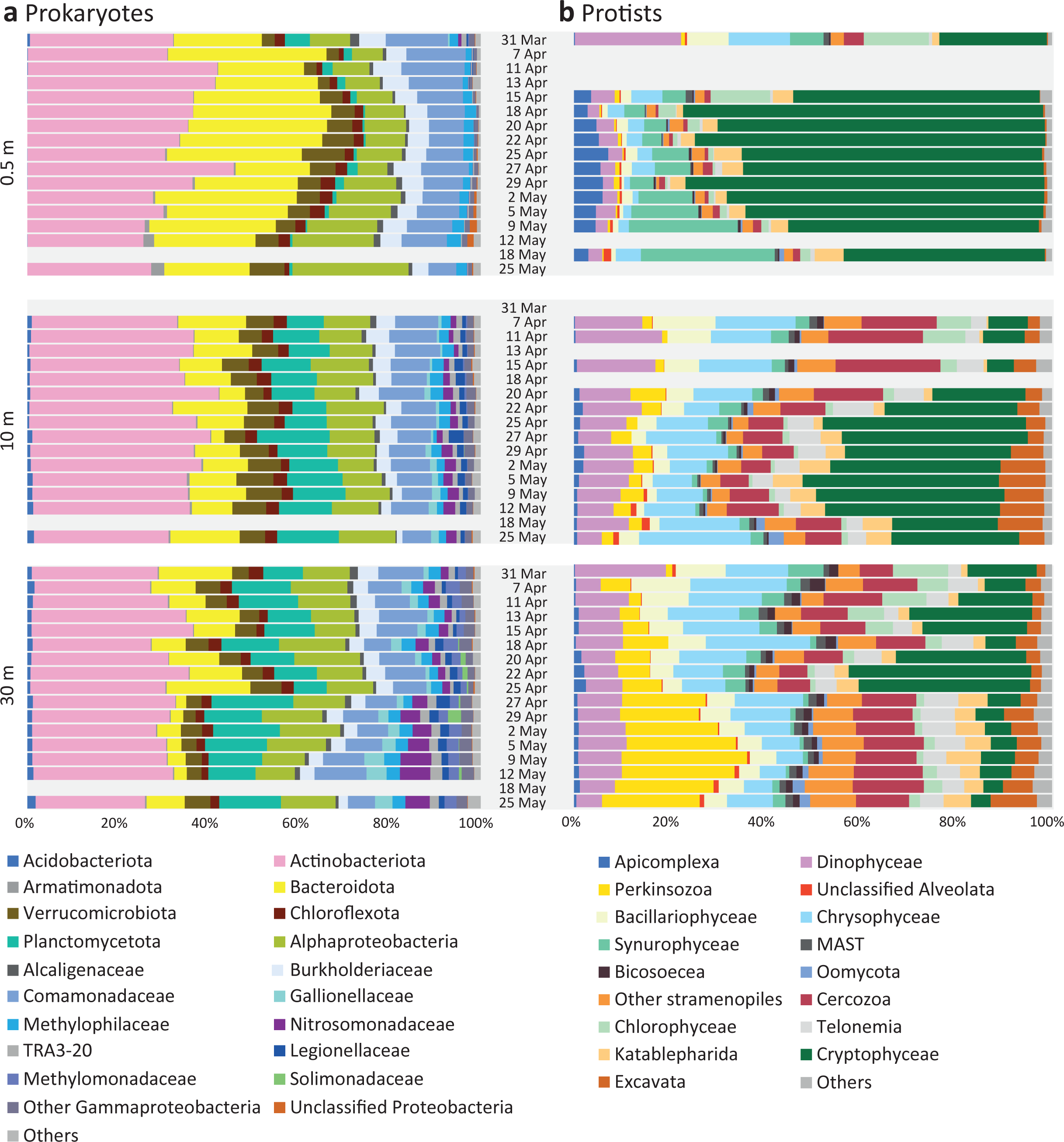
Prokaryotic and protistan community composition at three depths of Římov reservoir during the study. The gaps indicate missing samples. a – prokaryotic community. The majority of groups are resolved at phylum to class levels, with the exception of Gammaproteobacteria where several families were shown due to high heterogeneity in their distribution. b – protistian community. The groups are resolved at phylum to class level, with the exception of Supergroup Excavata, which was dominated by kinetoplastea. Ciliophora were excluded from the analysis of protists due to homogeneous distribution with high proportions in all the three layers. The figure including ciliophora is available in **Additional file 6**.

The epilimnion was characterised by a high contribution of Bacteroidota (max. 35%) and increasing read counts of Alphaproteobacteria (max. 26%) and Armatimonadota (max. 3%) towards the study end (**Figure 4a**). The eukaryotic community was dominated by cryptophytes with the highest contribution of 70% on 18^th^ April (**Figure 4b**). Synurophytes also contributed a significant number of reads, especially towards the study end (up to 30%). Apicomplexa, almost absent at the beginning, increased in relative abundances as the study progressed and reached 10% on 25^th^ April.

Among prokaryotes in the metalimnion, a high proportion of Planctomycetota was present throughout the study period, in contrast to negligible counts in the upper water layer. Moreover, populations of Acidobacteriota, Gallionellaceae, Nitrosomonadaceae, Legionellaceae and TRA3-20 established in the metalimnion albeit at low percentages (**Figure 4a**). Similar to the epilimnion, cryptophytes were the most dominant eukaryotic group and represented up to 40% on 25^th^ April and 5^th^ May (**Figure 4b**). In contrast, Dinophyta, Chrysophyta, Cercozoa, Telonema and Excavata had significantly higher contributions to the metalimnetic community.

In the hypolimnion, we observed a collapse of Bacteroidota population after 25^th^ April, the date corresponding to the breakpoint clearly separating the meta- from hypolimnion according to nMDS analysis (**Figure 3**). While the majority of bacterial groups were common to both metalimnetic and hypolimnetic samples (**Figure 4a**), Methylomonadaceae and Solimonadaceae were present exclusively in the hypolimnion. The eukaryotic community in the hypolimnion was dominated by Perkinsozoan sequences (max. 30%), especially towards the study end (**Figure 4b**). Similar to Bacteroidota, cryptophyte sequences drastically dropped after 25^th^ April which reflected the separation of hypolimnion from the metalimnion (**Figure 3, 4b**). Chrysophytes had a relatively high contribution in the hypolimnion during the early phase of the campaign but dropped considerably towards the study end. Cercozoa, Telonemia and Dinophyta had relatively high and stable proportions throughout the campaign.

### Dynamics of important groups of microbial eukaryotes in the water column

Abundances of eight eukaryotic groups were quantified by CARD-FISH using specific probes (**Table 1**, **Figure 5, Additional files 5, 7**). Cryptophytes dominated in the epilimnion for about 3 weeks from late April to early May (max. 4.3×10^3^ cells ml^-1^) and decreased towards the study end. Their abundances in the meta– and hypolimnion remained relatively low (< 1.4×10^3^ cells ml^-1^). Epilimnetic abundances of the aplastidic CRY1 lineage of cryptophytes initially increased to 1.6×10^3^ cells ml^-1^ (34% of the total eukaryotes) on April 13^th^ (**Figure 5**) and decreased to 0.4×10^3^ cells ml^-1^ towards the study end. In the metalimnion, abundances of this lineage showed similar but less pronounced dynamics and reached maxima on 20^th^ April with 1.1×10^3^ cells ml^-1^ (27 % of total eukaryotes). Their abundance in the hypolimnion was relatively low (< 0.6×10^3^ cells ml^-1^, < 22% of total eukaryotes). Katablepharids targeted by probe Kat-1492 were not very abundant in the epilimnion with exception of two peaks of ca. 0.2×10^3^ cells ml^-1^ (**Figure 5**). In the metalimnion, their abundance gradually increased and represented up to 7% of total eukaryotes. In the hypolimnion, this lineage showed low numbers throughout the study period, in contrast to the katablepharids detected with probe Kat2-651 which were found exclusively in the hypolimnion with up to 0.3×10^3^ cells ml^-1^ **(Figure 5**).

**Figure 5:**
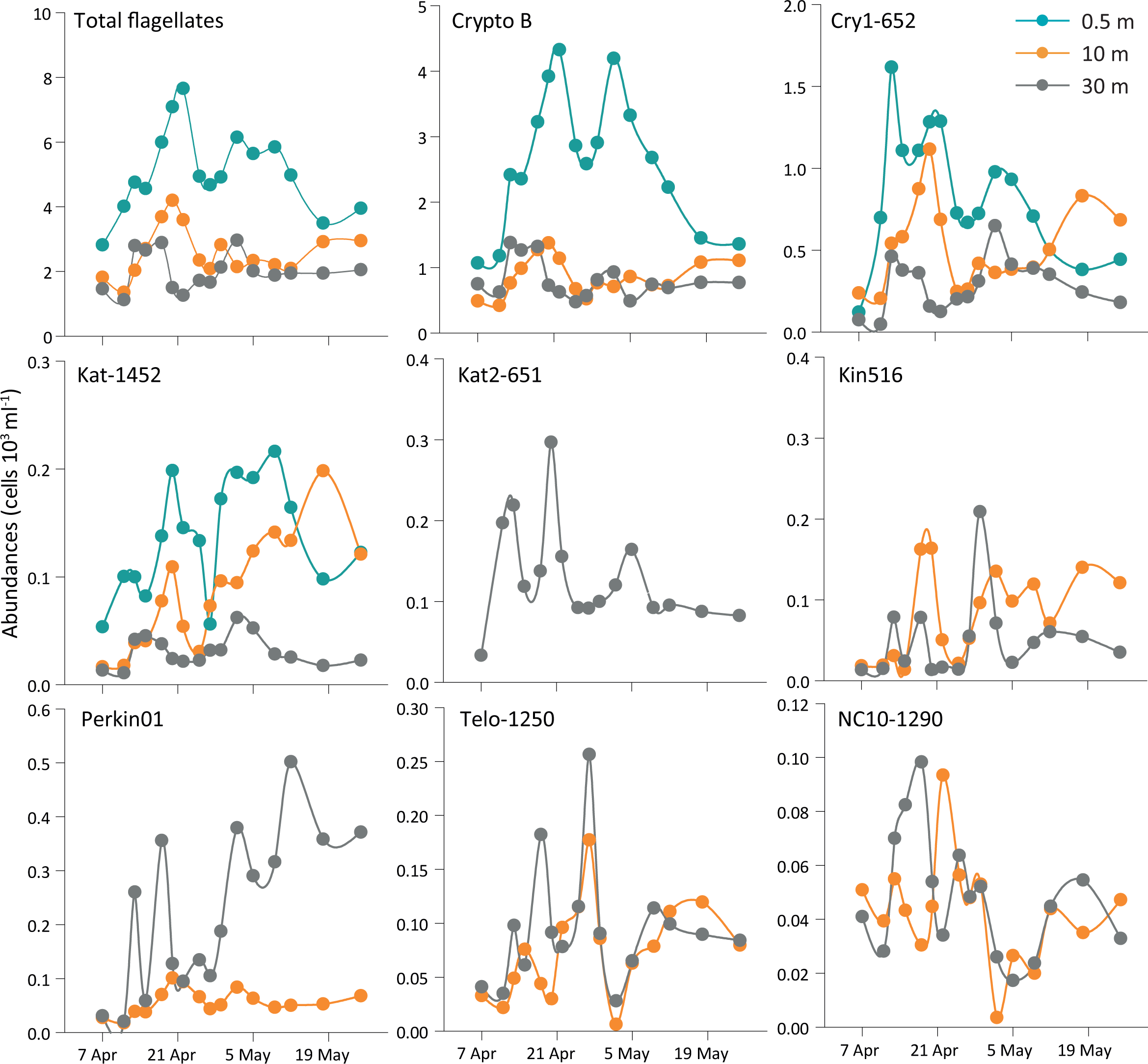
Absolute abundances of particular flagellate groups at three depths of Římov reservoir obtained by CARD-FISH. Left to right from top-total eukaryotes by DAPI staining, cryptophytes targeted by probe Crypto B, CRY1 lineage of cryptophytes targeted by probe Cry1-652, katablepharids targeted by probe Kat-1452, katablepharid clade 2 targeted by probe Kat2-651, kinetoplastids targeted by probe Kin516, Perkinsozoa clade 1 targeted by probe Perkin01, Telonemids targeted by probe Telo-1250, and Cercozoa Novel Clade 10 targeted by probe NC10-1290. A figure representing relative abundances of flagellates can be found in the supplemental material (**Additional file 7**).

Kinetoplastida, Perkinsozoa, Telonema and Cercozoa Novel Clade 10 were detected only in the meta- and hypolimnion (**Figure 5**). In the metalimnion, Kinetoplastida increased with some oscillations till the study end with maxima of 0.18×10^3^ cells ml^-1^. Similarly, their abundances fluctuated considerably with maxima of 0.21×10^3^ cells ml^-1^ in the hypolimnion. Perkinsozoans targeted by probe Perkin01 had relatively low abundances in the metalimnion (max. 0.1×10^3^ cells ml^-1^), while they reached up to 0.5×10^3^ cells ml^-1^ in the hypolimnion (**Figure 5**). Telonema were abundant in meta- and hypolimnion samples with the highest peaks recorded on 25^th^ April with 0.17×10^3^ cells ml^-1^ and 0.26×10^3^ cells ml^-1^, respectively. The Novel Clade 10 of Cercozoa showed similar abundance patterns to those observed for Telonema (**Figure 5**). However, maximal abundances were 2-3 times lower, i.e., 0.09×10^3^ cells ml^-1^ and 0.10×10^3^ cells ml^-1^ in meta- and hypolimnion, respectively.

## Discussion

High-frequency sampling, adhering tightly to the typical doubling time of microbes, allowed us to follow the community assembly at three depths during the transition from mixis to stratification in the water column. Interestingly, both protistan and prokaryotic community development showed strikingly similar dynamics in the different water strata of the reservoir (**Figure 3**). The eukaryotic and bacterial communities in the epilimnion gradually diverged from those in the meta- and hypolimnion soon after the mixing, responding to the increase in temperature and light intensity. Moreover, towards the end of the campaign when the water column became stratified, we observed a clear separation between meta- and hypolimnetic communities.

### Formation of water strata-associated communities and detected microbial interactions

We performed a network analysis to examine the connections between and within the eukaryotic and bacterial communities **(Figure 6, Additional file 8**). The majority of potential interactions were located in two large clusters consisting of three modules each. The modules of the smaller cluster reflected the temporal development within the epilimnion. Phototrophic eukaryotes, containing typical members of spring blooms[31] such as *Cryptomonas* (Cryptophyta), *Chlamydomonas* (Chlorophyta) and Chrysophyta had central positions. They were excessively linked to members of Bacteroidota, namely Flavobacteriales, Chitinophagales and Sphingobacteriales, which contributed the highest proportion of nodes in this cluster. These bacterial groups and Sphingomonas (present in module one) were reported as efficient decomposers of phytoplankton derived polymers during spring blooms[28, 33, 34]. In modules two and three, which represent the later phase of the study, the potential consumers of small molecular substances such as *Polynucleobacter* sp., Actinobacteriota and copiotrophic Comamonadaceae were present. These bacteria are well known members of spring bloom and disturbance succession[27–29, 35]. Ciliates present in the epilimnetic network cluster are known for their substratum-attached lifestyle, e.g., fine-filter feeding *Vorticella* sp.[65, 66] that were connected to colonial chrysophytes, or are moderately efficient bacterial grazers such as *Rimostrombidium* and *Strombidimum* spp.[62], which were extensively connected to bacterial nodes.

**Figure 6:**
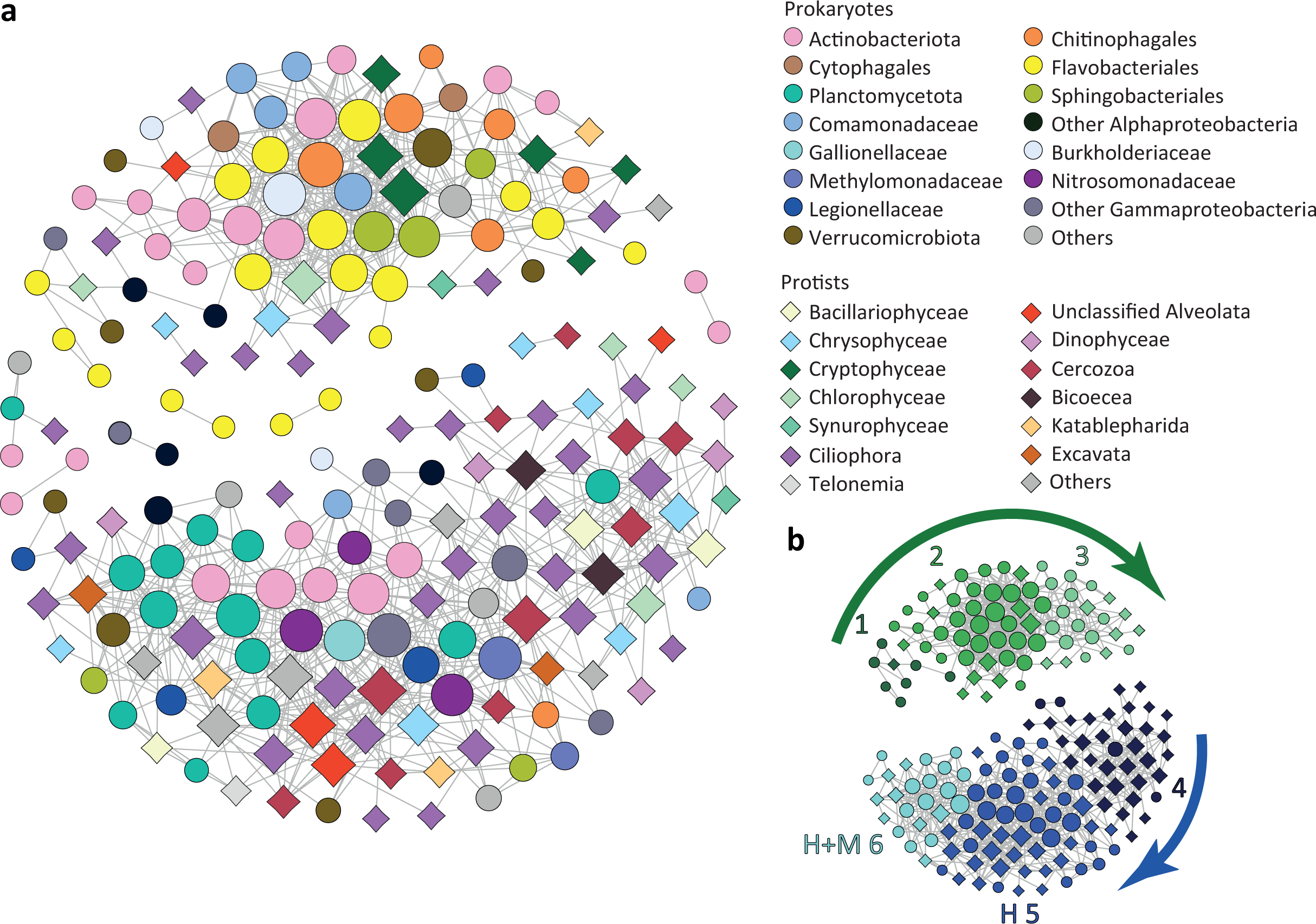
Network analysis based on the most abundant ASVs from protistan and prokaryotic communities. a – network: upper cluster represents the community dominating in the epilimnion, lower cluster represents the community dominating in the hypolimnion. Prokaryotic nodes are displayed as circles, protistan nodes as diamonds. b – the main modules detected in the network. The arrows indicate directions of temporal shifts between modules. Differentiation between modules H 5 and H+M 6 is based on spatial parameters as module 6 was also well represented by members of the metalimnion community.

The modules in the large meta- and hypolimnion associated cluster showed a limited temporal resolution. The module four of the large meta- and hypolimnion associated cluster (**Figure 6, Additional file 8**) represented the earliest part of the sampling period and consisted mainly of eukaryotes with sessile lifestyles. It was organized around centric diatoms which bloomed in the reservoir at the beginning of sampling (**Figure 2a**). A highly interconnected bacterial node in this part of the network was affiliated with Pirellulaceae (Planctomycetota), which were previously recorded in Římov reservoir during algal blooms and found attached to particles[67]. This Pirellulaceae node is linked to algal species and bacterivorous eukaryotes such as diatoms, cercozoans and ciliates. Chrysophytes and dinophytes were additional algal groups detected in module four, which are known to survive in low-light conditions and have mixotrophic or heterotrophic representatives[3, 8, 17]. Ciliophora comprised the lion share of this module and were represented mainly by organisms with sessile lifestyle (*Vorticella, Tokophrya,* etc.) or loricate forms with psychrophilic preferences (*Tintinnidium*)[68]. Interestingly, four *Legionella* sp. ASVs were part of the hypolimnion network. They are often reported as symbionts vs parasites of unicellular eukaryotes[69, 70] and were linked to eukaryotic ASVs in our analysis. Moreover, in two cases this was a selective connection to a single eukaryote (ciliate and cercozoa) and simultaneous association with Verrucomicrobiota (**Figure 6, Additional file 8**). Among flagellates, the heterotrophic groups bicosoecea and cercozoa had a significant presence in module four. Bicosoecids are characterized as mainly attached flagellates feeding on bacteria or more rarely free-living and feeding on particle-associated bacteria[8, 71]. Cercozoans are also widely reported from freshwater environments[16, 23] and were shown to feed on bacteria and small eukaryotes[7, 72] or to parasitize phytoplankton[23, 73]. Both strategies corroborate the associations detected in the co-occurrence network (**Figure 6**). Modules five and six (**Figure 6, Additional file 8**), predominantly consisting of bacteria, appear to be associated with lake snow (macroscopic organic aggregates)[74]. Rather than showing a temporal distinction, these modules exhibit a depth-related separation. Module five was more aligned with the hypolimnion, while module six represent both meta- and hypolimnion samples. Microbes from module six may possess fast colonization potential of lake snow aggregates that sink through the water column. Planctomycetota prevalent in this network section were mainly represented by members of Nemodikiaceae (CL500-3 group), known for their psychrophilic nature and particle-attached lifestyle[67, 75]. Planctomycetes are capable of peptide degradation through the so-called ‘planctosome’ complex bound to the outer membrane[67, 76]. Such putative micro-environments high in aminopeptidase activity, rich in labile amino acids[77] and ammonia could also benefit free-living Nanopelagicales (Actinobacteriota), specialists in the uptake of amino acids[35, 78], which were dominant in module five. Ammonia oxidizers such as Nitrosomonadaceae were also present in this module and were connected to nitrite oxidizers, e.g. *Nitrotoga* sp. (Gallionellaceae). Nitrate produced by these bacteria was shown to positively influence the methane oxidation efficiency of *Methylobacter* spp. (Methylomonadaceae)[79]. ASVs affiliated with *Methylobacter* were linked to *Nitrotoga* sp. and Nitrosomonadaceae in the network. Similar to module four, the microbial eukaryotes present here, i.e., excavates and cercozoans are known to be substrate-attached or associated with detritus and are mainly reported from hypolimnia of freshwater lakes[24, 80]. Different feeding strategies could be responsible for lineage-specific distribution of protists in the water column, as one lineage of katablepharids (Kat2-651) was detected with CARD-FISH exclusively in the hypolimnion, while another lineage was abundant in the epi- and metalimnion (**Figure 5**). Omnivorous and predatory strains of katablepharids are described in the literature[7, 38, 81]. Excavates (Kinetoplastea and Diplonema) are typically found in the hypolimnion of freshwater lakes during summer[22, 24, 80], or in hypertrophic, shallow lakes rich in suspended organic particles[38]. In our study, the presence of kinetoplastids in the metalimnion and hypolimnion was confirmed by both sequencing and CARD-FISH data. This group represents two nodes in the network modules five and six associated with the meta- and hypolimnion (**Figure 6, Additional file 8**), in line with reports of kinetoplastids feeding on bacteria associated with detritus particles[8, 82] sinking from the epilimnion to deeper strata at the end of the spring bloom[80]. One additional node in module five was represented by Telonema. This group has been reported from a wide range of freshwater habitats[18, 20, 24, 83]. However, our study is the first to track their population dynamics using a specific oligonucleotide probe (**Table 1, Additional file 5**), revealing that telonemids are almost absent in surface waters and mainly inhabit the deep-water layers (**Figures 4, 5**).

### Dynamics of dominant protistan groups

Cryptophyta was the most abundant eukaryotic group dominating in all epi- and metalimnion samples, as detected by both sequencing and CARD-FISH methods (**Figures 4, 5, Additional files 6, 7**). Microscopic observations showed the high abundances of big (10-30 µm long) chloroplast-bearing but also small aplastidic cryptophytes. Highly abundant heterotrophic CRY1 lineage[7, 38] accounted on average for 30% of the total cryptophytes targeted by the general Crypto B probe (**Figure 5**). This lineage did not show clear associations within the network but contributed substantially to the first maximum of HNF observed in epi- and metalimnion and probably played an important role as bacterivores in line with recent findings of high bacterial uptake rates of CRY1[38, 39]. The balance between auto- and heterotrophic cryptophytes during the springtime is related to the availability of ample sunlight, nutrients and prokaryotic prey[27, 39, 84].

Perkinsozoa dominated the deep-water communities (**Figures 4, 5**) and were not present in the co-occurrence network. Perkinsozoa comprise putative parasitic protists widely distributed in marine and freshwaters[21, 23, 85–87]. In our study, the contribution of Perkinsozoan ASVs and cell abundance increased with the onset of stratification (**Figures 4, 5**), reaching up to 25% of total ASVs and 26% of total eukaryotes in the hypolimnion, similar to previous studies[21, 23, 28]. With CARD-FISH analysis, perkinsozoans were observed both free-living and inside of free-living protists (**Additional file 5**). An additional taxon well represented in the sequencing data and absent in the network analysis was Apicomplexa, a poorly understood group especially in freshwater environments. Apicomplexa were reported as obligate intracellular parasites mainly affecting fish and phytoplankton[88]. In our data set, they were more abundant in the epilimnion and significantly correlated with cryptophytes (r^2^=0.600, p=2.719×10^-10^). Apicomplexa and perkinsozoa were not part of the network probably due to their putative parasitic association with higher eukaryotes that were not included in analyses.

### Microbial food web organization in spring

Algal blooms are the major factors shaping the spring microbial community not only in the epilimnion[89, 90] but also in the deeper strata, due to the enhanced particle flux from the surface waters[80, 91]. The bloom of chrysophytes, diatoms and large cryptophytes in the reservoir at the beginning of the sampling campaign was likely decimated by zooplankton or viral lysis[28]. However, ciliates dominated by raptorial prostomes such as *Urotricha* spp. and *Balanion planktonicum* also contributed to the phytoplankton reduction[64, 92] (**Figure 1d**). Cladocerans, especially large sized *Daphnia* spp. were highly abundant during the first two weeks of sampling and seemed to represent the main driver responsible for the decline of phytoplankton bloom in mid-April. The highest abundances of smaller grazers, i.e., Rotifera were observed close to the maxima of their corresponding favourite prey chrysophytes and cryptophytes (**Figure 2**). *Polyarthra* spp., which were shown to selectively feed on chrysophytes[93], were first in the succession and were followed by omnivorous *Keratella* sp. preferably feeding on chrysophytes and cryptophytes[94]. However, *Cyclops vicinus* seemed to drastically reduce rotifer population, similar to previous observations of spring plankton succession in Římov reservoir[95]. The simultaneous establishment of a stable population of *Eudiaptomus gracilis* probably did not contribute much to the rotifers’ top-down control, but this copepod successfully replaced rotifers and daphnids as a powerful algal grazer in the later phase[96]. *E. gracilis* was also shown to exhibit strong influence on the lower food web organization due to a high clearance rate of ciliates[97]. The drop in ciliate densities, including high proportions of bacterivorous or omnivorous species, most likely resulted in a short-term increase in bacterial numbers (**Figure 1d, f**). Notably, ciliates were almost equally important bacterivores than HNF during times of high ciliate abundances (**Additional file 9**). Ciliates, especially raptorial prostomes also contributed to the reduction of HNF in the epilimnion (**Figure 1d**, e). In addition, approximately half of the ciliate community during its peak abundance in early April was composed of typical flagellate hunters such as *Balanion planktonicus* and *Urotricha* spp. (data not shown)[7, 92, 98]. After the drop of ciliate abundance, the protistan bulk bacterivory rate was largely attributed to HNF (**Additional file 9**) dominated by aplastidic cryptophytes such as CRY1 lineage[38, 39] and omnivorous katablepharids (**Figure 5**)[7, 38].

Communities in the deeper strata showed a clear dominance of heterotrophic groups although we did not follow the dynamics of higher trophic levels due to their low abundances. However, the highly complex network between prokaryotes and protists in the hypolimnion (**Figure 6**) indicates a considerable increase of bacterivorous, parasitic, and detritivorous strategies in these strata.

## Conclusions

In this study, we followed the dynamics of organisms < 200 µm in three different water column layers of a freshwater reservoir at high-temporal resolution during spring. The results unveiled parallel community assembly patterns for protists and prokaryotes, revealing an early separation of epilimnetic communities and subsequent differentiation between the meta- and hypolimnetic layers. Besides confirming a prevalence of phototrophic and predatory strategies among epilimnetic protists, we observed the emergence of organisms affiliated with Perkinsozoa, Telonemia, Kinetoplastida, and Cercozoa in the meta- and hypolimnion, indicating a dominance of particle-associated lifestyles and parasitic and detritivorous strategies in deeper strata during the spring period. Furthermore, we showcased diverse associations between bacterial and protistan taxa, ranging from substrate degradation-related to parasitic. These associations followed temporal successions and displayed depth-specific dynamics. Sequence-based and microscopic techniques allowed for the integration of protists into a holistic picture of the complex community dynamics during springtime. Such a hybrid approach appears to be a powerful tool for integrating various groups of organisms in temporal and spatial dynamics, enhancing our comprehension of microbial interactions and the functioning of freshwater ecosystems.

## Declarations

### Ethics approval and consent to participate

Not applicable.

### Consent for publication

Not applicable.

### Availability of data and material

The sequence data generated from the 16S and 18S rRNA gene amplicon sequencing was submitted to the European Nucleotide Archive (ENA) and are available under the BioProject: PRJEB66298, [https://www.ebi.ac.uk/ena/browser/view/PRJEB66298].

### Author’s Contributions

IM, VG and KŠ conceived the study. IM and TS wrote the manuscript with the input from all authors. IM performed CARD-FISH, data analysis and interpretation. VG performed sampling, CARD-FISH and data analysis. TS performed sampling, sequence data analysis and interpretation, and design of figures. MS performed sampling, phylogenetic analyses and construction of CARD-FISH probes. KŠ performed microscopic counts, bacterivory rates estimation, data analysis and interpretation. PZ, PR, JS and MD performed sampling and sample analysis. All the authors contributed to critical revisions and approved the final version of the manuscript.

## Acknowledgements

We are thankful to Radka Malá for excellent laboratory assistance.

## Additional materials

**File name** – Additional file 1

**File format** – .pdf

**Title and description of data** – Chemistry data. DOC - dissolved organic carbon, DN - dissolved nitrogen, DSi - dissolved silica, TP - total phosphorus, DP - dissolved phosphorus, DRP dissolved reactive phosphorus, A254-400 absorbance measured at corresponding wavelength (nm).

**File name** – Additional file 2

**File format** – .pdf

**Title and description of data** – Randomized axelerated maximum likelihood (RAxML) tree (100 bootstraps) of katablepharids. Branches with bootstrap support <20% were multifurcated, probe targets are marked by different colors. Asterisks indicate sequences not targeted by probes.

**File name** – Additional file 3

**File format** – .pdf

**Title and description of data** – Randomized axelerated maximum likelihood (RAxML) tree (100 bootstraps) of Cercozoa including Novel Clade 10 (NC10). Branches with bootstrap support <20% were multifurcated, probe targets are marked by different colors. Asterisks indicate sequences not targeted by probe.

**File name** – Additional file 4

**File format** – .pdf

**Title and description of data** – Randomized axelerated maximum likelihood (RAxML) tree (100 bootstraps) of Telonema. Branches with bootstrap support <20% were multifurcated, probe targets are marked by different colors. Asterisks indicate sequences not targeted by the probes, # indicates sequence that is too short to be checked for the target region.

**File name** – Additional file 5

**File format** – .pdf

**Title and description of data** – Microphotographs displaying: a. different lineages of protists hybridized with CARD-FISH probes designed for this study (Kat2-651, Telo-1250 and NC10-1290), cell hybridized with Kat-1452 shown for comparison; b. different lifestyles observed for Perkinsozoa hybridized with Perkin01 (upper row free-living, lower row protist-associated). The scale bar applies for all images. Microphotographs were produced using Zeiss Imager Z2, Carl Zeiss, Oberkochen, DE equipped with a Colibri LED system and the following filter sets: DAPI 49 (Excitation 365; Beamsplitter TFT 395; Emission BP 445/50), fluorescein 38 HE (Excitation BP 470/40; Beamsplitter TFT 495; Emission BP 525/50).

**File name** – Additional file 6

**File format** – .pdf

**Title and description of data** – Protistan community composition at three depths of Římov reservoir during the study. The gaps indicate missing samples. The groups are resolved at phylum to class level, with the exception of Supergroup Excavata, which was dominated by kinetoplastea.

**File name** – Additional file 7

**File format** – .pdf

**Title and description of data** – Relative abundances of particular flagellate groups in three depths of Římov reservoir obtained with CARD-FISH analysis. Left to right from top- cryptophytes targeted with Crypto B probe, CRY1 lineage of cryptophytes targeted with Cry1-652 probe, katablepharids targeted with Kat-1452 probe, katablepharid clade 2 targeted with Kat2-651 probe, kinetoplastids targeted with Kin516 probe, Perkinsozoa clade 1 targeted with Perkin01 probe, Telonemids targeted with Telo-1250 probe, and Cercozoa novel clade 10 targeted with NC10-1290 probe.

**File name** – Additional file 8

**File format** – .pdf

**Title and description of data** – Network analysis based on the most abundant ASVs from protistan and prokaryotic communities. a – network: upper cluster represents the community dominating in the epilimnion, lower cluster represents the community dominating in the hypolimnion. Prokaryotic nodes are displayed as circles, protistan nodes as diamonds. b – the main modules detected in the network. The arrows indicate directions of temporal shifts between modules. Differentiation between modules H 5 and H+M 6 is based on spatial parameters as members of module 6 were better resented in the metalimnion communities. Prokaryotic and protistan nodes are organized into modules and listed below, accompanied by heatmaps based on Z scores calculated for each module. Samples are grouped according to water column layers, with the time course depicted from left to right.

**File name** – Additional file 9

**File format** – .pdf

**Title and description of data** – Total grazing by protists in the epilimnion

