## Additional File 1 for "Depth-dependent dynamics of protist communities as an integral part of spring succession in a freshwater reservoir"

**Additional file 1:** Chemistry data. DOC - dissolved organic carbon, DN - dissolved nitrogen, DSi - dissolved silica, TP - total phosphorus, DP - dissolved phosphorus, DRP dissolved reactive phosphorus, A254-400 absorbance measured at corresponding wavelength (nm).

**Epilimnion 0.5m**

| Date | pH | DOC mg/L | DN mg/L | DSi mg/L | TP mg/L | DP µg/l | DRP µg/l | $NH_4$-N µg/l | $NO_3$-N mg/L | A254 | A300 | A350 | A400 |
|---|---|---|---|---|---|---|---|---|---|---|---|---|---|
| 31-Mar-16 | 7.07 | 4.69 | 1.35 | 6.7 | 33.0 | 20.4 | 4.8 | 8.3 | 1.21 | 0.160 | 0.087 | 0.038 | 0.016 |
| 07-Apr-16 | 7.44 | 4.79 | 1.41 | 5.7 | 25.6 | 10.3 | 2.1 | 4.3 | 1.08 | 0.156 | 0.085 | 0.037 | 0.015 |
| 11-Apr-16 | 7.32 | 4.94 | - | 5.7 | 20.1 | 11.2 | 2.9 | 31.8 | 1.11 | 0.155 | 0.083 | 0.036 | 0.015 |
| 13-Apr-16 | 7.38 | 4.64 | 1.51 | 5.6 | 22.8 | 10.5 | 3.0 | 28.0 | 1.07 | 0.145 | 0.073 | 0.030 | 0.009 |
| 15-Apr-16 | 7.38 | 4.98 | 1.6 | 5.6 | 20.6 | 10.1 | 3.1 | 32.2 | 1.11 | 0.159 | 0.086 | 0.039 | 0.017 |
| 18-Apr-16 | 7.29 | 5.07 | 1.54 | 5.9 | 20.5 | 12.5 | 2.8 | 26.8 | 1.16 | 0.154 | 0.082 | 0.035 | 0.014 |
| 20-Apr-16 | 7.45 | 4.71 | 1.32 | 5.2 | 14.6 | 9.0 | 2.8 | 19.4 | 1.09 | 0.152 | 0.081 | 0.035 | 0.014 |
| 22-Apr-16 | 7.65 | 4.71 | 1.43 | 5.4 | 15.4 | 10.6 | 3.9 | 24.0 | 1.08 | 0.153 | 0.082 | 0.035 | 0.015 |
| 25-Apr-16 | 7.46 | 4.81 | 1.43 | 5.3 | 14.6 | 9.1 | 2.0 | 13.0 | 1.10 | 0.151 | 0.081 | 0.034 | 0.014 |
| 27-Apr-16 | 7.45 | 4.62 | 1.44 | 5.2 | 17.3 | 9.0 | 6.0 | 16.0 | 1.09 | 0.148 | 0.079 | 0.034 | 0.014 |
| 29-Apr-16 | 7.57 | 4.68 | 1.44 | 2.6 | 17.8 | 9.5 | 3.8 | 28.0 | 1.08 | 0.149 | 0.079 | 0.034 | 0.014 |
| 02-May-16 | 7.35 | 5.32 | 1.74 | 5.3 | 17.1 | 10.9 | 3.5 | 38.0 | 1.07 | 0.149 | 0.080 | 0.033 | 0.013 |
| 05-May-16 | 7.11 | 4.51 | 1.56 | 5.3 | 15.8 | 8.1 | 1.6 | 13.0 | 1.03 | 0.145 | 0.077 | 0.033 | 0.013 |
| 09-May-16 | 7.69 | 4.53 | 1.54 | 5.5 | 16.4 | 8.9 | 1.7 | 16.0 | 0.99 | 0.143 | 0.075 | 0.031 | 0.013 |
| 12-May-16 | 7.52 | 4.42 | 1.51 | 5.2 | 14.4 | 9.1 | 3.3 | 3.0 | 0.91 | 0.138 | 0.074 | 0.031 | 0.012 |
| 18-May-16 | 7.72 | 4.40 | 1.39 | 4.7 | 17.9 | 9.1 | 1.3 | 32.7 | 0.93 | 0.139 | 0.072 | 0.030 | 0.012 |

**Hypolimnion 30m**

| Date | pH | DOC mg/L | DN mg/L | DSi mg/L | TP mg/L | DP µg/l | DRP µg/l | $NH_4$-N µg/l | $NO_3$-N mg/L | A254 | A300 | A350 | A400 |
|---|---|---|---|---|---|---|---|---|---|---|---|---|---|
| 31-Mar-16 | 7.08 | 4.75 | 1.31 | 6.51 | 29.0 | 19.5 | 9.8 | 17.2 | 1.14 | 0.162 | 0.088 | 0.039 | 0.016 |
| 07-Apr-16 | 7.09 | 4.75 | 1.37 | 5.77 | 26.3 | 18.0 | 11.6 | 2.4 | 1.06 | 0.161 | 0.087 | 0.038 | 0.015 |
| 11-Apr-16 | 7.14 | 4.73 | - | 5.79 | 26.6 | 17.3 | 9.0 | 14.0 | 1.11 | 0.159 | 0.086 | 0.038 | 0.015 |
| 13-Apr-16 | 7.13 | 4.71 | 1.49 | 5.77 | 23.5 | 15.4 | 9.3 | 17.1 | 1.08 | 0.148 | 0.075 | 0.031 | 0.010 |
| 15-Apr-16 | 7.34 | 4.96 | 1.58 | 6.09 | 23.4 | 16.5 | 9.2 | 14.6 | 1.09 | 0.161 | 0.088 | 0.040 | 0.017 |
| 18-Apr-16 | 7.12 | 5.07 | 1.51 | 5.85 | 25.1 | 19.0 | 12.0 | 7.9 | 1.17 | 0.160 | 0.087 | 0.038 | 0.015 |
| 20-Apr-16 | 7.11 | 4.71 | 1.31 | 5.61 | 23.7 | 16.9 | 12.1 | 8.1 | 1.07 | 0.159 | 0.086 | 0.038 | 0.015 |
| 22-Apr-16 | 7.31 | 4.78 | 1.44 | 5.79 | 20.5 | 15.1 | 10.3 | 6.0 | 1.14 | 0.158 | 0.085 | 0.037 | 0.015 |
| 25-Apr-16 | 7.38 | 4.75 | 1.43 | 5.64 | 20.9 | 14.8 | 9.7 | 14.0 | 1.12 | 0.157 | 0.085 | 0.037 | 0.015 |
| 27-Apr-16 | 7.20 | 4.83 | 1.46 | 5.58 | 23.4 | 18.3 | 14.6 | 15.0 | 1.12 | 0.158 | 0.085 | 0.037 | 0.015 |
| 29-Apr-16 | 7.27 | 4.76 | 1.46 | 3.32 | 23.7 | 17.4 | 12.5 | 15.0 | 1.10 | 0.158 | 0.086 | 0.037 | 0.015 |
| 02-May-16 | 7.02 | 4.69 | 1.57 | 5.78 | 24.3 | 17.9 | 12.6 | 17.0 | 1.12 | 0.158 | 0.085 | 0.037 | 0.015 |
| 05-May-16 | 7.03 | 4.72 | 1.55 | 5.31 | 22.8 | 17.0 | 11.5 | 9.0 | 1.09 | 0.157 | 0.085 | 0.037 | 0.015 |
| 09-May-16 | 7.14 | 4.59 | 1.55 | 6.22 | 24.6 | 17.4 | 11.6 | 2.0 | 1.09 | 0.157 | 0.085 | 0.037 | 0.015 |
| 12-May-16 | 7.05 | 4.54 | 1.59 | 5.85 | 23.4 | 16.9 | 12.7 | 0.0 | 1.10 | 0.156 | 0.086 | 0.037 | 0.015 |
| 18-May-16 | 7.16 | 4.59 | 1.49 | 5.65 | 24.4 | 18.9 | 10.8 | 14.4 | 1.06 | 0.155 | 0.084 | 0.036 | 0.014 |
| 25-May-16 | 6.99 | 4.51 | - | 5.76 | 23.5 | 16.2 | 12.1 | 18.0 | 1.07 | 0.152 | 0.084 | 0.036 | 0.014 |
