## Additional File 2 for "Depth-dependent dynamics of protist communities as an integral part of spring succession in a freshwater reservoir"

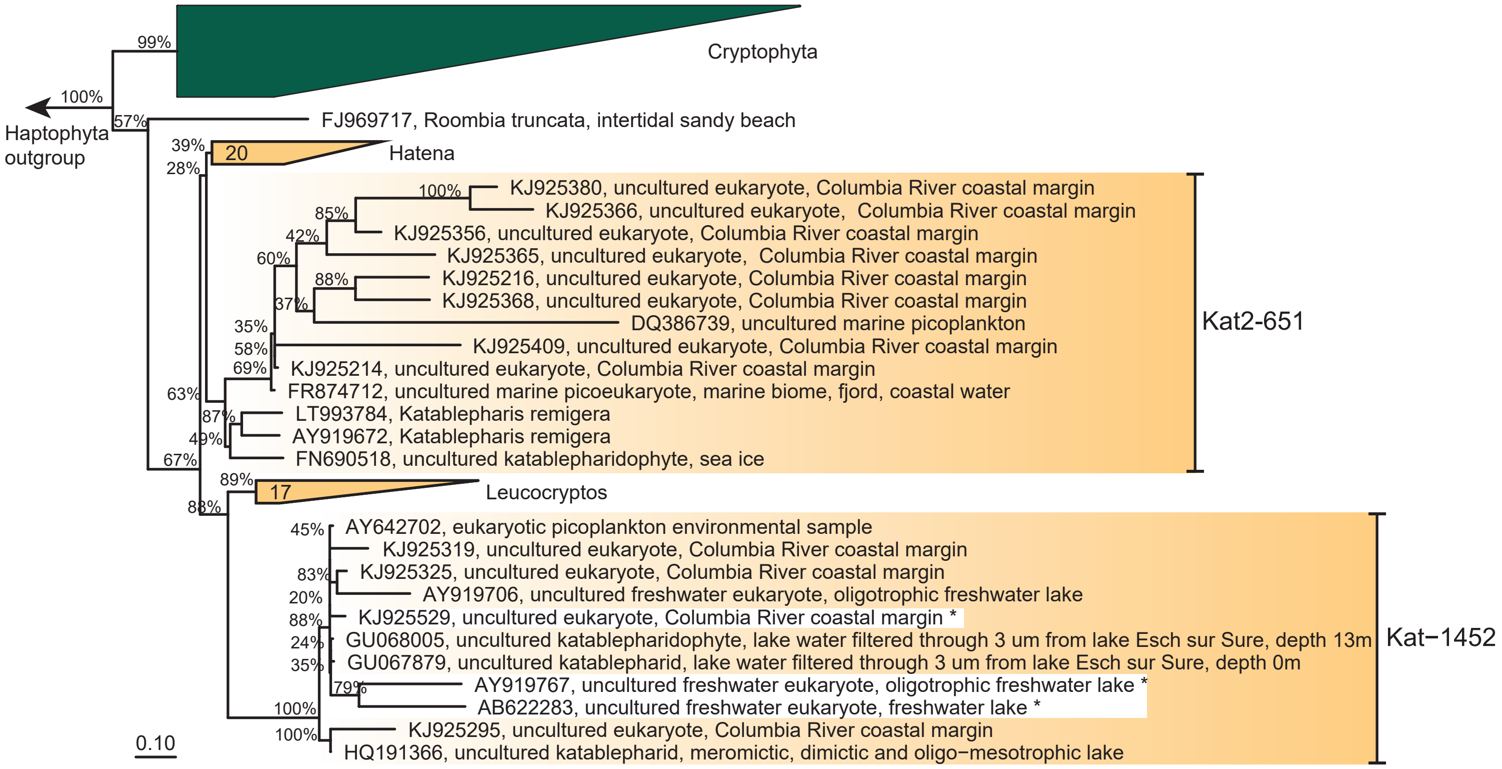


**Additional file 2:** Randomized axelerated maximum likelihood (RAxML) tree (100 bootstraps) of katablepharids. Branches with bootstrap support <20% were multifurcated, probe targets are marked by different colors. Asterisks indicate sequences not targeted by probes.
