## Additional File 5 for "Depth-dependent dynamics of protist communities as an integral part of spring succession in a freshwater reservoir"

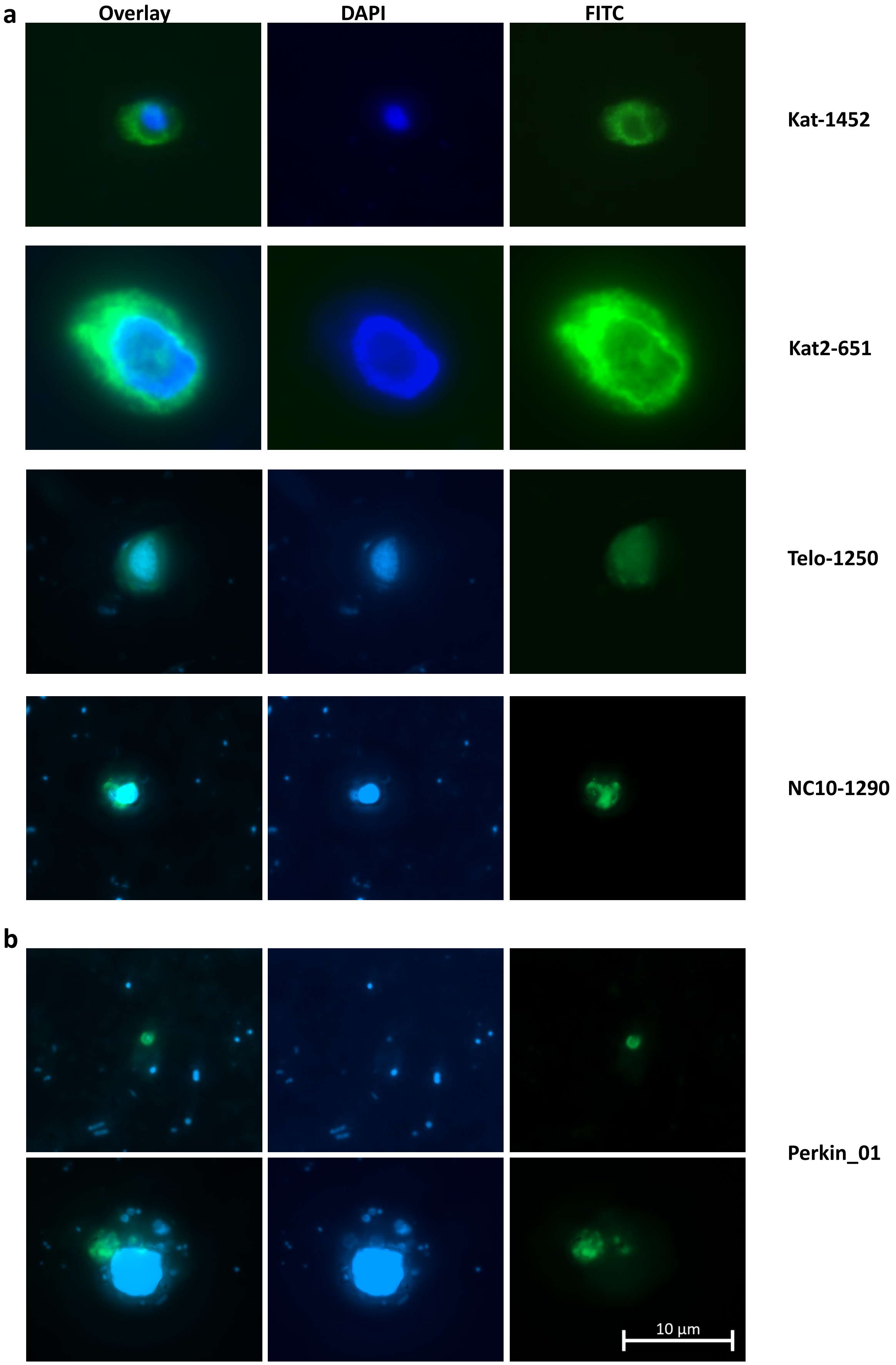


**Additional file 5:** Microphotographs displaying: **a.** different lineages of protists hybridized with CARD-FISH probes designed for this study (Kat2-651, Telo-1250 and NC10-1290), cell hybridized with Kat-1452 shown for comparison; **b.** different lifestyles observed for Perkinsozoa hybridized with Perkin_01 (upper row free-living, lower row protist-associated). The scale bar applies for all images.

Microphotographs were produced using Zeiss Imager Z2, Carl Zeiss, Oberkochen, DE equipped with a Colibri LED system and the following filter sets: DAPI 49 (Excitation 365; Beamsplitter TFT 395; Emission BP 445/50), fluorescein 38 HE (Excitation BP 470/40; Beamsplitter TFT 495; Emission BP 525/50).
