## Additional File 6 for "Depth-dependent dynamics of protist communities as an integral part of spring succession in a freshwater reservoir"

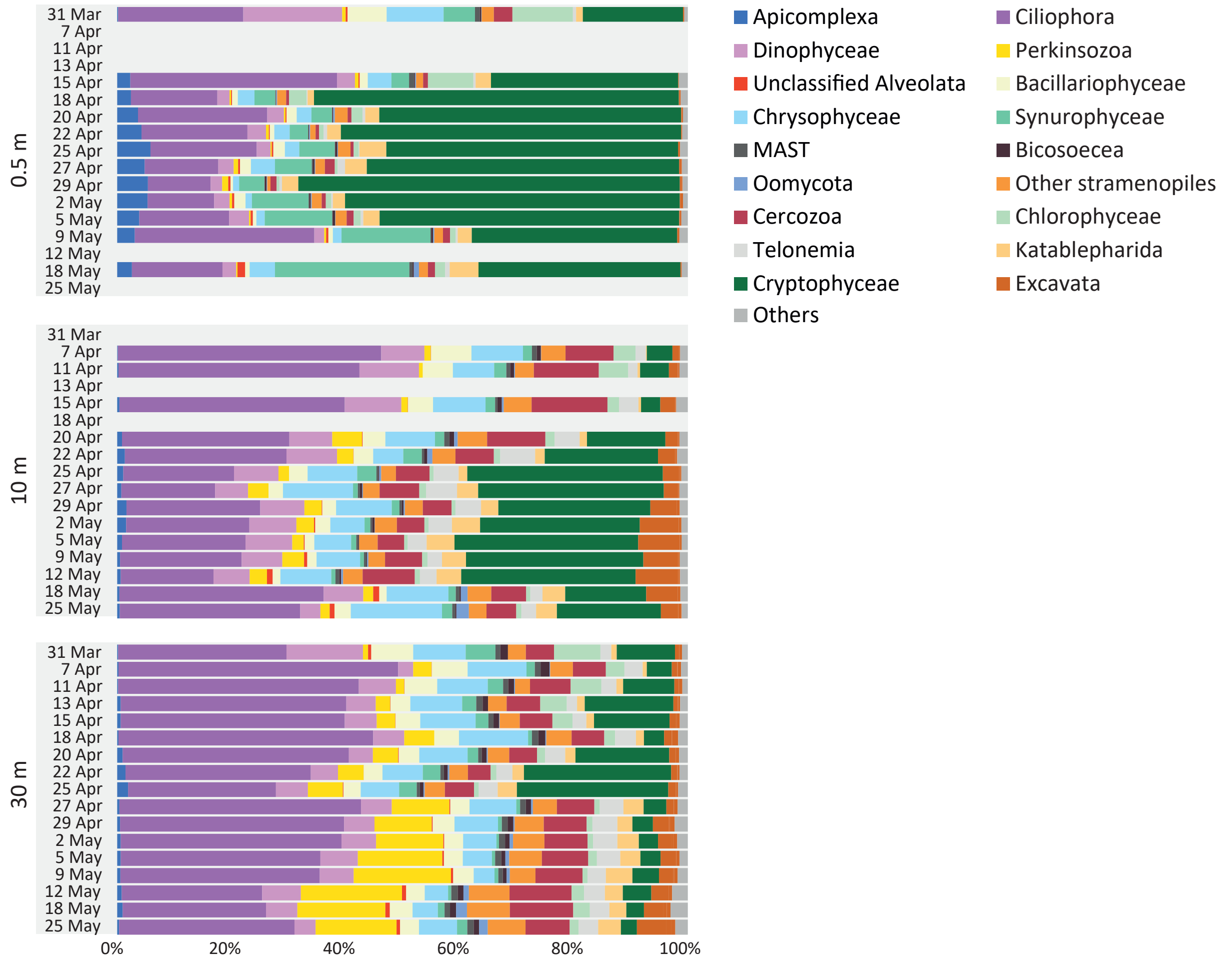


**Additional file 6:** Protistan community composition at three depths of Římov reservoir during the study. The gaps indicate missing samples. The groups are resolved at phylum to class level, with the exception of Supergroup Excavata, which was dominated by kinetoplastea.
