## Additional File 7 for "Depth-dependent dynamics of protist communities as an integral part of spring succession in a freshwater reservoir"

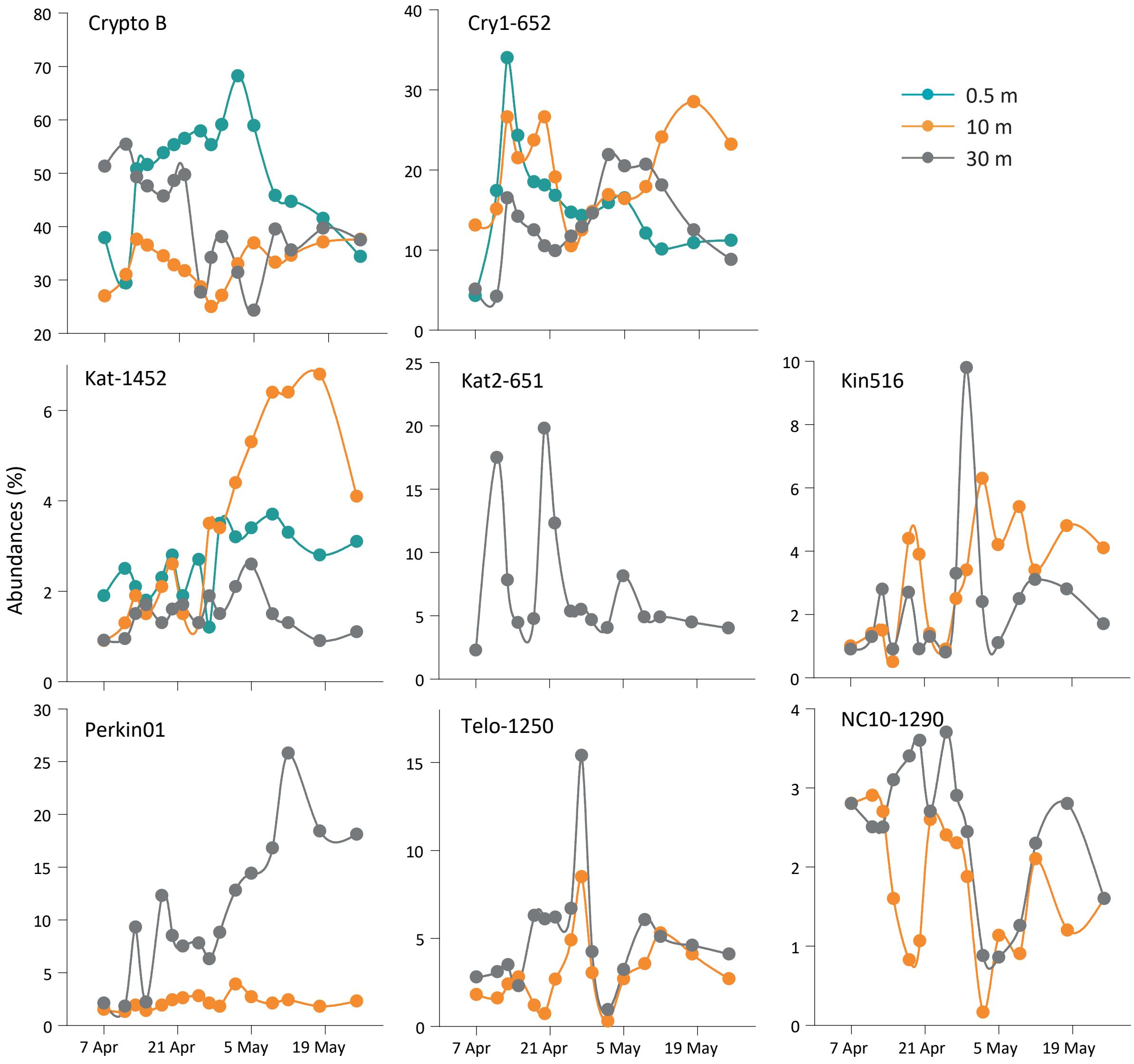


**Additional file 7:** Relative abundances of particular flagellate groups in three depths of Římov reservoir obtained with CARD-FISH analysis. Left to right from top- cryptophytes targeted with Crypto B probe, CRY1 lineage of cryptophytes targeted with Cry1-652 probe, katablepharids targeted with Kat-1452 probe, katablepharid clade 2 targeted with Kat2-651 probe, kinetoplastids targeted with Kin516 probe, Perkinsozoa clade 1 targeted with Perkin01 probe, Telonemids targeted with Telo-1250 probe, and Cercozoa novel clade 10 targeted with NC10-1290 probe.
