## Additional File 8 for "Depth-dependent dynamics of protist communities as an integral part of spring succession in a freshwater reservoir"

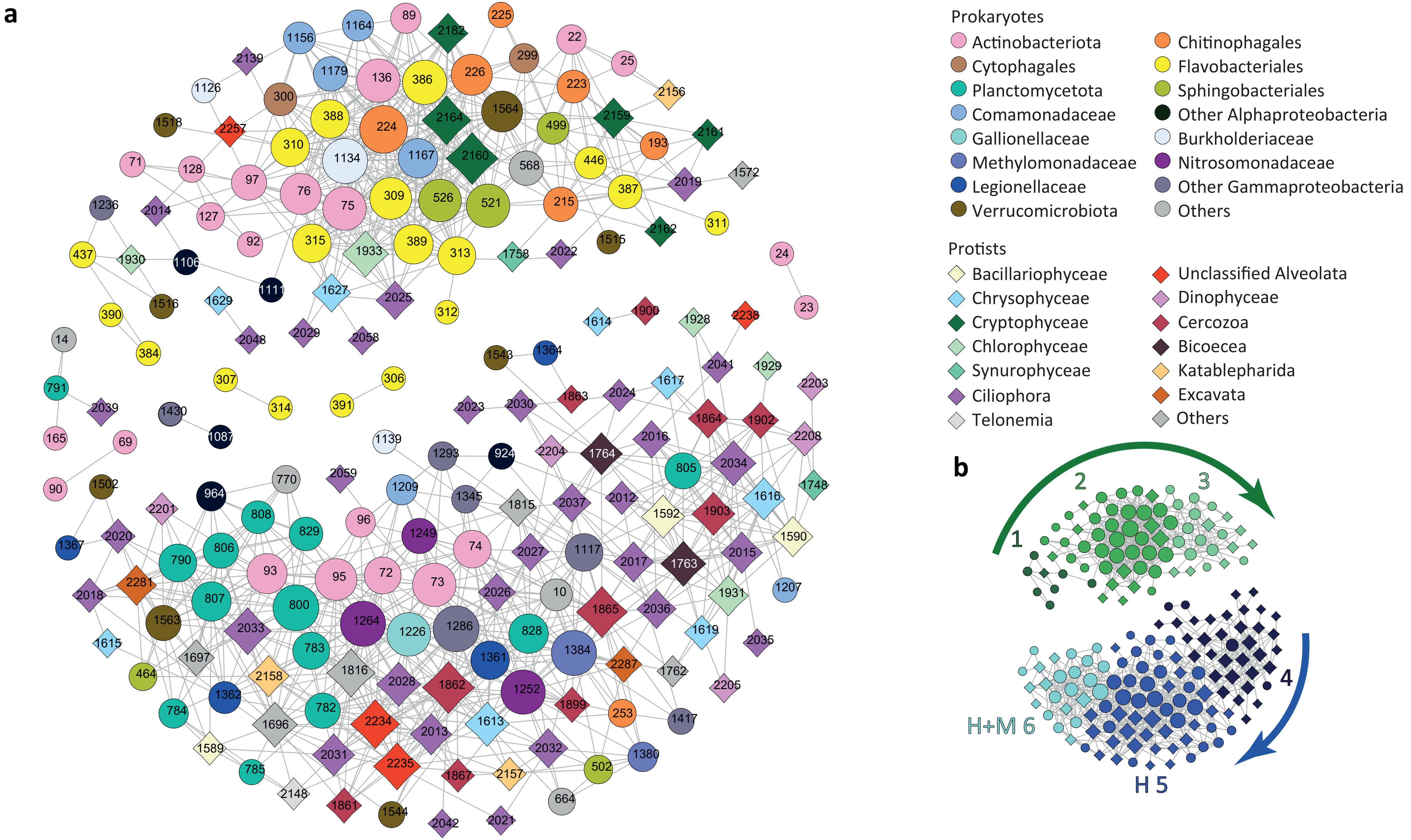


**Additional file 8:**

Network analysis based on the most abundant ASVs from protistan and prokaryotic communities. **a** – network: upper cluster represents the community dominating in the epilimnion, lower cluster represents the community dominating in the hypolimnion. Prokaryotic nodes are displayed as circles, protistan nodes as diamonds. **b** – the main modules detected in the network. The arrows indicate directions of temporal shifts between modules. Differentiation between modules H 5 and H+M 6 is based on spatial parameters as members of module 6 were better resented in the metalimnion communities. Prokaryotic and protistan nodes are organized into modules and listed below, accompanied by heatmaps based on Z scores calculated for each module. Samples are grouped according to water column layers, with the time course depicted from left to right.

### Module 1

| Phylum | Class | Order | Family | Genus | ID |
|---|---|---|---|---|---|
| Bacteroidota | Bacteroidia | Flavobacteriales | Flavobacteriaceae | Flavobacterium | 384 |
| Bacteroidota | Bacteroidia | Flavobacteriales | Flavobacteriaceae | Flavobacterium | 390 |
| Bacteroidota | Bacteroidia | Flavobacteriales | Flavobacteriaceae | Flavobacterium | 437 |
| Verrucomicrobiota | Verrucomicrobiae | NA | NA | NA | 1516 |
| Proteobacteria | Gammaproteobacteria | Burkholderiales | Methylophilaceae | Methylotenera | 1236 |
| Proteobacteria | Alphaproteobacteria | Sphingomonadales | Sphingomonadaceae | Sphingorhabdus | 1106 |
| Proteobacteria | Alphaproteobacteria | Sphingomonadales | Sphingomonadaceae | Sphingorhabdus | 1111 |
| Archaeplastida | Chlorophyta | Chlorophyceae | Chlamydomonadales | Chlamydomonadales_X | 1930 |

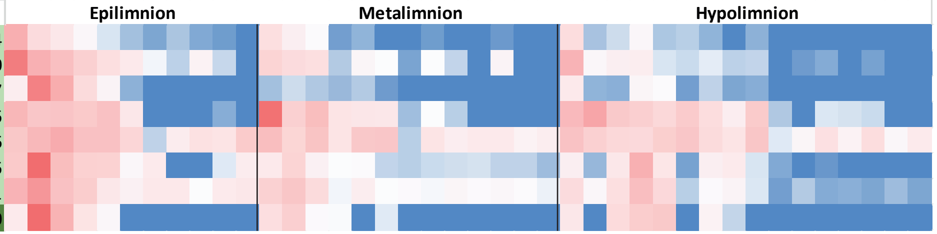


### Module 2

| Phylum | Class | Order | Family | Genus | ID |
|---|---|---|---|---|---|
| Actinobacteriota | Actinobacteria | Frankiales | Sporichthyaceae | Candidatus Planktophila | 71 |
| Actinobacteriota | Actinobacteria | Frankiales | Sporichthyaceae | Candidatus Planktophila | 75 |
| Actinobacteriota | Actinobacteria | Frankiales | Sporichthyaceae | Candidatus Planktophila | 76 |
| Actinobacteriota | Actinobacteria | Frankiales | Sporichthyaceae | hgcI clade | 89 |
| Actinobacteriota | Actinobacteria | Frankiales | Sporichthyaceae | hgcI clade | 92 |
| Actinobacteriota | Actinobacteria | Frankiales | Sporichthyaceae | hgcI clade | 97 |
| Actinobacteriota | Actinobacteria | Frankiales | Sporichthyaceae | NA | 127 |
| Actinobacteriota | Actinobacteria | Frankiales | Sporichthyaceae | NA | 128 |
| Actinobacteriota | Actinobacteria | Micrococcales | Microbacteriaceae | Candidatus Limnoluna | 136 |
| Bacteroidota | Bacteroidia | Chitinophagales | Chitinophagaceae | Sediminibacterium | 224 |
| Bacteroidota | Bacteroidia | Chitinophagales | Chitinophagaceae | Sediminibacterium | 225 |
| Bacteroidota | Bacteroidia | Chitinophagales | Chitinophagaceae | Sediminibacterium | 226 |
| Bacteroidota | Bacteroidia | Cytophagales | Spirosomaceae | Pseudarcicella | 299 |
| Bacteroidota | Bacteroidia | Cytophagales | Spirosomaceae | Pseudarcicella | 300 |
| Bacteroidota | Bacteroidia | Flavobacteriales | Crocinitomicaceae | Fluviicola | 309 |
| Bacteroidota | Bacteroidia | Flavobacteriales | Crocinitomicaceae | Fluviicola | 310 |
| Bacteroidota | Bacteroidia | Flavobacteriales | Crocinitomicaceae | Fluviicola | 312 |
| Bacteroidota | Bacteroidia | Flavobacteriales | Crocinitomicaceae | Fluviicola | 313 |
| Bacteroidota | Bacteroidia | Flavobacteriales | Crocinitomicaceae | Fluviicola | 315 |
| Bacteroidota | Bacteroidia | Flavobacteriales | Flavobacteriaceae | Flavobacterium | 386 |
| Bacteroidota | Bacteroidia | Flavobacteriales | Flavobacteriaceae | Flavobacterium | 387 |
| Bacteroidota | Bacteroidia | Flavobacteriales | Flavobacteriaceae | Flavobacterium | 388 |
| Bacteroidota | Bacteroidia | Flavobacteriales | Flavobacteriaceae | Flavobacterium | 389 |
| Bacteroidota | Bacteroidia | Sphingobacteriales | Sphingobacteriaceae | Pedobacter | 521 |
| Bacteroidota | Bacteroidia | Sphingobacteriales | Sphingobacteriaceae | Solitalea | 526 |
| Proteobacteria | Gammaproteobacteria | Burkholderiales | Burkholderiaceae | Polynucleobacter | 1126 |
| Proteobacteria | Gammaproteobacteria | Burkholderiales | Burkholderiaceae | Polynucleobacter | 1134 |
| Proteobacteria | Gammaproteobacteria | Burkholderiales | Comamonadaceae | Acidovorax | 1156 |
| Proteobacteria | Gammaproteobacteria | Burkholderiales | Comamonadaceae | Limnohabitans | 1164 |
| Proteobacteria | Gammaproteobacteria | Burkholderiales | Comamonadaceae | Limnohabitans | 1167 |
| Proteobacteria | Gammaproteobacteria | Burkholderiales | Comamonadaceae | NA | 1179 |
| Verrucomicrobiota | Verrucomicrobiae | NA | NA | NA | 1518 |
| Verrucomicrobiota | Verrucomicrobiae | Verrucomicrobiales | Verrucomicrobiaceae | NA | 1564 |
| Alveolata | Ciliophora | Spirotrichea | Strombidiida | Pelagostrombidiidae | 2014 |
| Hacrobia | Haptophyta | Prymnesiophyceae | Prymnesiales | Chrysochromulinaceae | 2257 |
| Alveolata | Ciliophora | NA | NA | NA | 2139 |
| Stramenopiles | Ochrophyta | Chrysophyceae | Chrysophyceae_X | NA | 1627 |
| Alveolata | Ciliophora | Oligohymenophorea | Peritrichia_2 | Sessilida | 2058 |
| Alveolata | Ciliophora | Spirotrichea | Tintinnida | TIN_03 | 2029 |
| Alveolata | Ciliophora | Spirotrichea | Hypotrichia | Oxytrichidae | 2025 |
| Hacrobia | Cryptophyta | Cryptophyceae | Cryptomonadales | Cryptomonadales_X | 2164 |
| Hacrobia | Cryptophyta | NA | NA | NA | 2182 |
| Archaeplastida | Chlorophyta | Chlorophyceae | Chlamydomonadales | Chlamydomonadales_X | 1933 |
| Hacrobia | Cryptophyta | Cryptophyceae | Cryptomonadales | Cryptomonadales_X | 2160 |

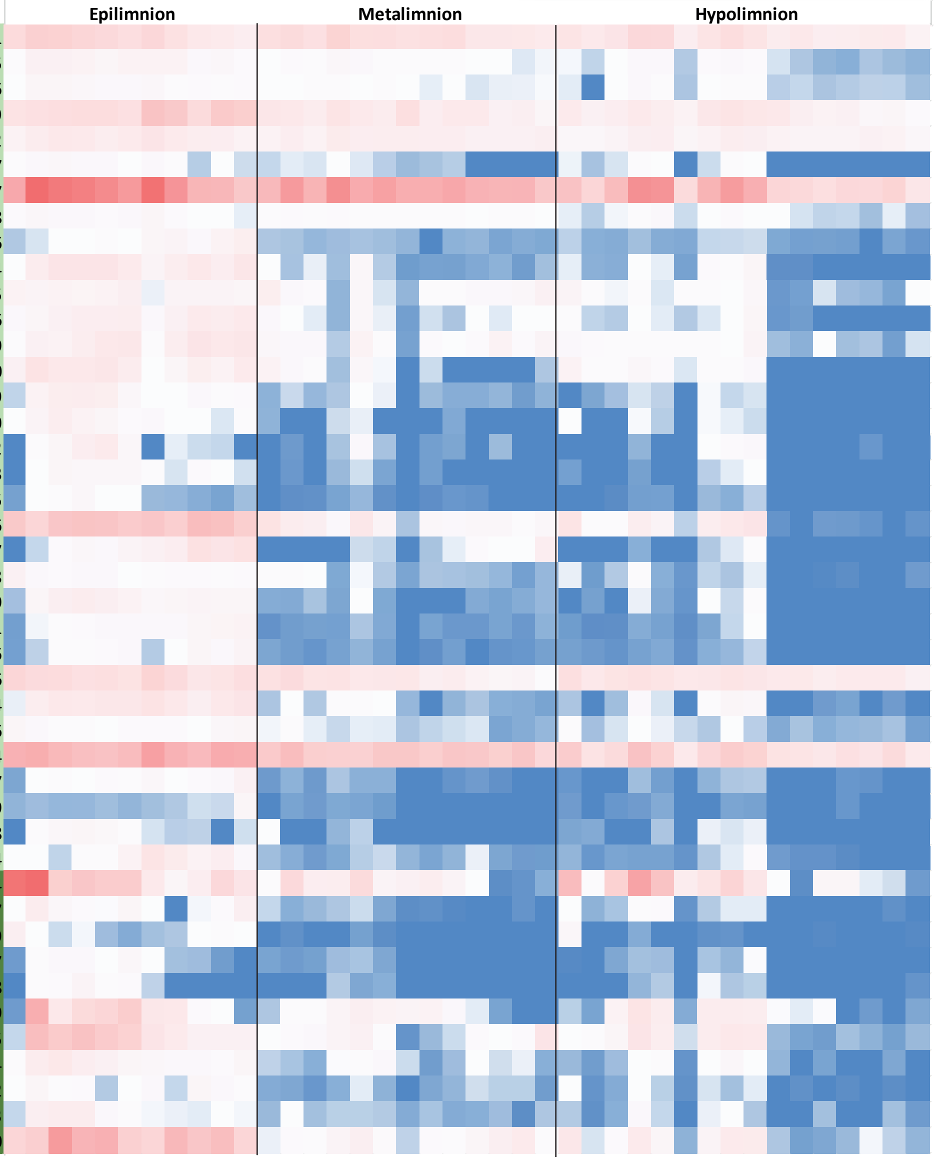


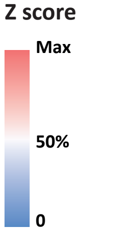


Prokaryotic ASVs

Protistan ASVs

### Module 3

| Phylum | Class | Order | Family | Genus | ID |
|---|---|---|---|---|---|
| Chloroflexi | SL56 marine group | NA | NA | NA | 568 |
| Bacteroidota | Bacteroidia | Chitinophagales | Chitinophagaceae | NA | 215 |
| Verrucomicrobiota | Verrucomicrobiae | NA | NA | NA | 1515 |
| Bacteroidota | Bacteroidia | Flavobacteriales | Crocinitomicaceae | Fluviicola | 311 |
| Bacteroidota | Bacteroidia | Flavobacteriales | NS9 marine group | NA | 446 |
| Bacteroidota | Bacteroidia | Chitinophagales | Chitinophagaceae | Edaphobaculum | 193 |
| Bacteroidota | Bacteroidia | Sphingobacteriales | NS11-12 marine group | NA | 499 |
| Bacteroidota | Bacteroidia | Chitinophagales | Chitinophagaceae | Sediminibacterium | 223 |
| Actinobacteriota | Acidimicrobiia | Microtrichales | Ilumatobacteraceae | CL500-29 marine group | 22 |
| Actinobacteriota | Acidimicrobiia | Microtrichales | Ilumatobacteraceae | CL500-29 marine group | 25 |
| Alveolata | Apicomplexa | Coccidiomorphea | Adeleida | NA | 1572 |
| Alveolata | Ciliophora | Spirotrichea | Strombidiida_A | Strombidiida_A_X | 2022 |
| Hacrobia | Katablepharidophyta | Katablepharidaceae | Katablepharidales | Katablepharidales_X | 2156 |
| Hacrobia | Cryptophyta | Cryptophyceae | Cryptomonadales | Cryptomonadales_X | 2159 |
| Hacrobia | Cryptophyta | Cryptophyceae | Cryptomonadales | Cryptomonadales_X | 2161 |
| Hacrobia | Cryptophyta | Cryptophyceae | Cryptomonadales | Cryptomonadales_X | 2162 |
| Stramenopiles | Ochrophyta | Synurophyceae | Synurales | Synurales_X | 1758 |
| Alveolata | Ciliophora | Spirotrichea | Choreotrichida | Strobilidiidae_D | 2019 |

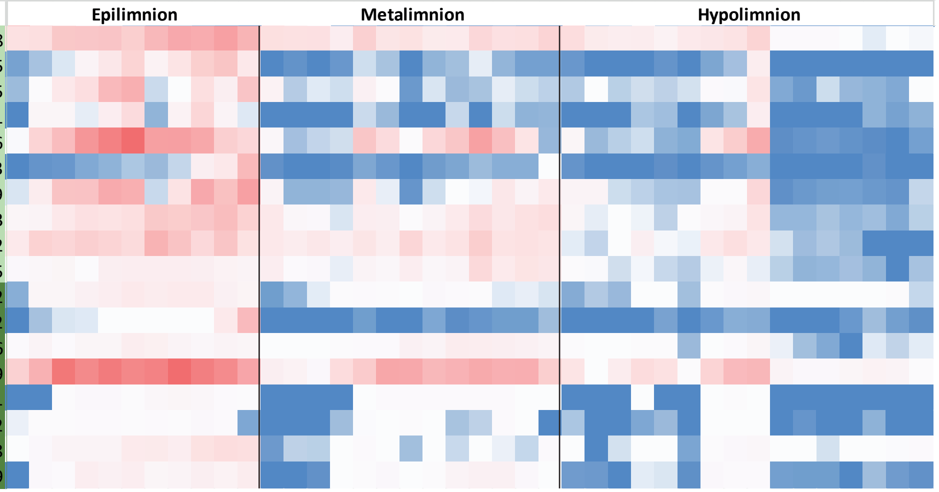


### Module 4

| Phylum | Class | Order | Family | Genus | ID |
|---|---|---|---|---|---|
| Planctomycetota | Planctomycetes | Pirellulales | Pirellulaceae | NA | 805 |
| Verrucomicrobiota | Verrucomicrobiae | Pedosphaerales | Pedosphaeraceae | NA | 1543 |
| Proteobacteria | Gammaproteobacteria | Legionellales | Legionellaceae | Legionella | 1364 |
| Proteobacteria | Gammaproteobacteria | Burkholderiales | Comamonadaceae | Polaromonas | 1207 |
| Stramenopiles | Ochrophyta | Bacillariophyta | Bacillariophyta_X | Polar-centric-Mediophyceae | 1590 |
| Stramenopiles | Ochrophyta | Bacillariophyta | Bacillariophyta_X | Polar-centric-Mediophyceae | 1592 |
| Stramenopiles | Ochrophyta | Chrysophyceae | Chrysophyceae_X | Chrysophyceae_Clade-C | 1616 |
| Stramenopiles | Ochrophyta | Chrysophyceae | Chrysophyceae_X | NA | 1617 |
| Stramenopiles | Ochrophyta | Chrysophyceae | Chrysophyceae_X | Chrysophyceae_Clade-E | 1619 |
| Stramenopiles | Ochrophyta | Synurophyceae | Synurales | Synurales_X | 1748 |
| Stramenopiles | Opalozoa | Bicoecea | Bicoecales | Bicoecaceae | 1763 |
| Stramenopiles | Opalozoa | Bicoecea | Bicoecales | Bicoecaceae | 1764 |
| Rhizaria | Cercozoa | NA | NA | NA | 1863 |
| Rhizaria | Cercozoa | Filosa-Thecofilosea | Ebriida | Botuliformidae | 1864 |
| Rhizaria | Cercozoa | NA | NA | NA | 1902 |
| Rhizaria | Cercozoa | NA | NA | NA | 1903 |
| Archaeplastida | Chlorophyta | Chlorophyceae | Chlamydomonadales | Chlamydomonadales_X | 1928 |
| Archaeplastida | Chlorophyta | Chlorophyceae | Chlamydomonadales | Chlamydomonadales_X | 1929 |
| Archaeplastida | Chlorophyta | Chlorophyceae | Chlamydomonadales | Chlamydomonadales_X | 1931 |
| Alveolata | Ciliophora | Oligohymenophorea | Scuticociliatia_2 | Histiobalantiidae | 2012 |
| Alveolata | Ciliophora | Spirotrichea | Tintinnida | Tintinnidiidae | 2015 |
| Alveolata | Ciliophora | Oligohymenophorea | Peritrichia_2 | Sessilida | 2016 |
| Alveolata | Ciliophora | Spirotrichea | Hypotrichia | NA | 2017 |
| Alveolata | Ciliophora | Litostomatea | Litostomatea_X | Litostomatea_XX | 2023 |
| Alveolata | Ciliophora | Colpodea | Colpodea_X | NA | 2024 |
| Alveolata | Ciliophora | NA | NA | NA | 2030 |
| Alveolata | Ciliophora | Oligohymenophorea | Peritrichia_2 | Sessilida | 2034 |
| Alveolata | Ciliophora | Spirotrichea | Tintinnida | Tintinnidiidae | 2035 |
| Alveolata | Ciliophora | Litostomatea | NA | NA | 2036 |
| Alveolata | Ciliophora | Phyllopharyngea | Suctoria | Tokophryidae | 2037 |
| Alveolata | Ciliophora | Oligohymenophorea | Peritrichia_2 | Sessilida | 2041 |
| Alveolata | Dinoflagellata | Dinophyceae | Gymnodiniales | Gymnodiniaceae | 2203 |
| Alveolata | Dinoflagellata | Dinophyceae | Prorocentrales | Prorocentraceae | 2204 |
| Alveolata | Dinoflagellata | Dinophyceae | Prorocentrales | Prorocentraceae | 2205 |
| Alveolata | Dinoflagellata | Dinophyceae | NA | NA | 2208 |
| Alveolata | NA | NA | NA | NA | 2238 |

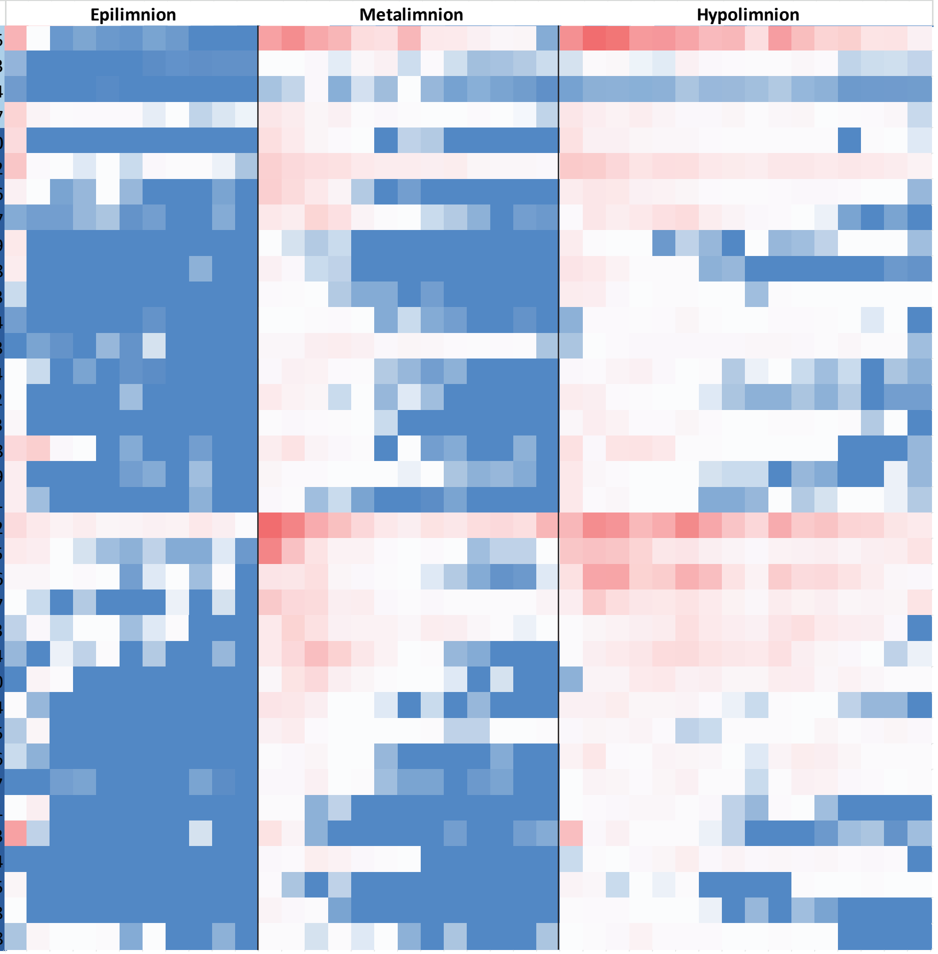


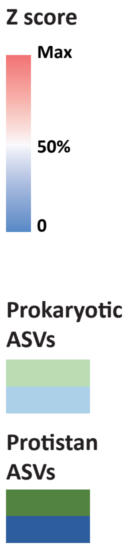

### Module 5

| Phylum | Class | Order | Family | Genus | ID |
|---|---|---|---|---|---|
| Acidobacteriota | Vicinamibacteria | Vicinamibacterales | NA | NA | 10 |
| Actinobacteriota | Actinobacteria | Frankiales | Sporichthyaceae | Candidatus Planktophila | 72 |
| Actinobacteriota | Actinobacteria | Frankiales | Sporichthyaceae | Candidatus Planktophila | 73 |
| Actinobacteriota | Actinobacteria | Frankiales | Sporichthyaceae | Candidatus Planktophila | 74 |
| Actinobacteriota | Actinobacteria | Frankiales | Sporichthyaceae | hgcI clade | 95 |
| Actinobacteriota | Actinobacteria | Frankiales | Sporichthyaceae | hgcI clade | 96 |
| Bacteroidota | Bacteroidia | Chitinophagales | NA | NA | 253 |
| Bacteroidota | Bacteroidia | Sphingobacteriales | NS11-12 marine group | NA | 502 |
| Myxococcota | Polyangia | mle1-27 | NA | NA | 664 |
| Planctomycetota | Phycisphaerae | Phycisphaerales | Phycisphaeraceae | CL500-3 | 782 |
| Planctomycetota | Phycisphaerae | Phycisphaerales | Phycisphaeraceae | CL500-3 | 783 |
| Planctomycetota | Phycisphaerae | Phycisphaerales | Phycisphaeraceae | CL500-3 | 785 |
| Planctomycetota | Planctomycetes | Pirellulales | Pirellulaceae | NA | 808 |
| Planctomycetota | Planctomycetes | Planctomycetales | Rubinisphaeraceae | SH-PL14 | 828 |
| Proteobacteria | Alphaproteobacteria | Rhizobiales | Beijerinckiaceae | alphaI cluster | 924 |
| Proteobacteria | Gammaproteobacteria | Burkholderiales | Alcaligenaceae | GKS98 freshwater group | 1117 |
| Proteobacteria | Gammaproteobacteria | Burkholderiales | Burkholderiaceae | Polynucleobacter | 1139 |
| Proteobacteria | Gammaproteobacteria | Burkholderiales | Comamonadaceae | Rhizobacter | 1209 |
| Proteobacteria | Gammaproteobacteria | Burkholderiales | Gallionellaceae | Candidatus Nitrotoga | 1226 |
| Proteobacteria | Gammaproteobacteria | Burkholderiales | Nitrosomonadaceae | GOUTA6 | 1249 |
| Proteobacteria | Gammaproteobacteria | Burkholderiales | Nitrosomonadaceae | NA | 1252 |
| Proteobacteria | Gammaproteobacteria | Burkholderiales | Nitrosomonadaceae | Nitrosospira | 1264 |
| Proteobacteria | Gammaproteobacteria | Burkholderiales | SC-I-84 | NA | 1286 |
| Proteobacteria | Gammaproteobacteria | Burkholderiales | TRA3-20 | NA | 1293 |
| Proteobacteria | Gammaproteobacteria | Diplorickettsiales | Diplorickettsiaceae | Rickettsiella | 1345 |
| Proteobacteria | Gammaproteobacteria | Legionellales | Legionellaceae | Legionella | 1361 |
| Proteobacteria | Gammaproteobacteria | Methylococcales | Methylomonadaceae | Methylobacter | 1380 |
| Proteobacteria | Gammaproteobacteria | Methylococcales | Methylomonadaceae | Methylobacter | 1384 |
| Proteobacteria | Gammaproteobacteria | Pseudomonadales | Pseudohongiellaceae | Blyi10 | 1417 |
| Verrucomicrobiota | Verrucomicrobiae | Pedosphaerales | Pedosphaeraceae | NA | 1544 |
| Verrucomicrobiota | Verrucomicrobiae | Verrucomicrobiales | Verrucomicrobiaceae | NA | 1563 |
| Stramenopiles | Ochrophyta | Chrysophyceae | Chrysophyceae_X | NA | 1613 |
| Stramenopiles | Ochrophyta | NA | NA | NA | 1696 |
| Stramenopiles | Pseudofungi | MAST-2 | MAST-2C | MAST-2C_X | 1762 |
| Stramenopiles | NA | NA | NA | NA | 1815 |
| Stramenopiles | NA | NA | NA | NA | 1816 |
| Rhizaria | Cercozoa | Filosa-Imbricatea | Filosa-Imbricatea_X | Novel-clade-2 | 1861 |
| Rhizaria | Cercozoa | Filosa-Imbricatea | Thaumatomonadida | Peregriniidae | 1862 |
| Rhizaria | Cercozoa | Filosa-Thecofilosea | Cryomonadida | Cryothecomonas-lineage | 1865 |
| Rhizaria | Cercozoa | Filosa-Thecofilosea | Cryomonadida | NA | 1867 |
| Rhizaria | Cercozoa | NA | NA | NA | 1899 |
| Alveolata | Ciliophora | NA | NA | NA | 2013 |
| Alveolata | Ciliophora | Oligohymenophorea | Hymenostomatia | Ophryoglenida | 2021 |
| Alveolata | Ciliophora | Spirotrichea | Choreotrichida | NA | 2026 |
| Alveolata | Ciliophora | NA | NA | NA | 2027 |
| Alveolata | Ciliophora | NA | NA | NA | 2028 |
| Alveolata | Ciliophora | NA | NA | NA | 2031 |
| Alveolata | Ciliophora | NA | NA | NA | 2032 |
| Alveolata | Ciliophora | Litostomatea | NA | NA | 2042 |
| Alveolata | Ciliophora | Ciliophora_X | Ciliophora_XX | Ciliophora_XXX | 2059 |
| Hacrobia | Telonemia | Telonemia_X | Telonemia_XX | Telonemia-Group-2 | 2148 |
| Hacrobia | Katablepharidophyta | Katablepharidaceae | Katablepharidales | Katablepharidales_X | 2157 |
| Hacrobia | Katablepharidophyta | Katablepharidaceae | Katablepharidales | Katablepharidales_X | 2158 |
| Alveolata | NA | NA | NA | NA | 2234 |
| Alveolata | NA | NA | NA | NA | 2235 |
| Excavata | Discoba | Kinetoplastea | Eubodonida | Bodonidae | 2287 |

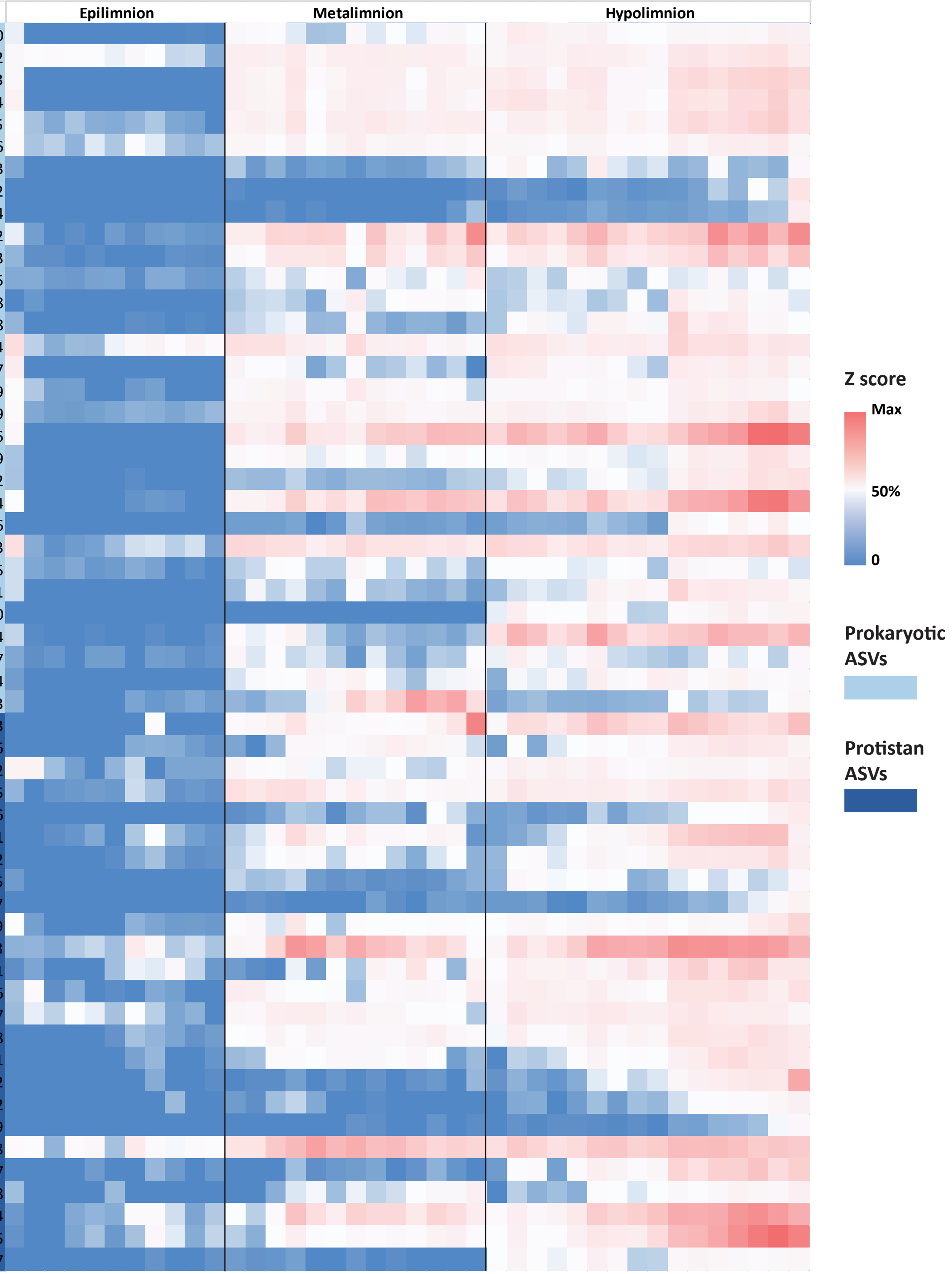

### Module 6

| Phylum | Class | Order | Family | Genus | ID |
|---|---|---|---|---|---|
| Planctomycetota | Planctomycetes | Gemmatales | Gemmataceae | NA | 790 |
| Planctomycetota | Planctomycetes | Isosphaerales | Isosphaeraceae | NA | 800 |
| Planctomycetota | Planctomycetes | Pirellulales | Pirellulaceae | NA | 806 |
| Planctomycetota | Planctomycetes | Pirellulales | Pirellulaceae | NA | 807 |
| Actinobacteriota | Actinobacteria | Frankiales | Sporichthyaceae | hgcI clade | 93 |
| Verrucomicrobiota | Verrucomicrobiae | Chthoniobacterales | Chthoniobacteraceae | Chthoniobacter | 1502 |
| Proteobacteria | Gammaproteobacteria | Legionellales | Legionellaceae | Legionella | 1367 |
| Bacteroidota | Bacteroidia | Sphingobacteriales | env.OPS 17 | NA | 464 |
| Planctomycetota | Phycisphaerae | Phycisphaerales | Phycisphaeraceae | CL500-3 | 784 |
| Proteobacteria | Alphaproteobacteria | Rhizobiales | Rhizobiales Incertae Se | NA | 964 |
| Patescibacteria | Saccharimonadia | Saccharimonadales | NA | NA | 770 |
| Planctomycetota | Planctomycetes | Planctomycetales | Schlesneriaceae | Schlesneria | 829 |
| Proteobacteria | Gammaproteobacteria | Legionellales | Legionellaceae | Legionella | 1362 |
| Stramenopiles | Ochrophyta | Bacillariophyta | Bacillariophyta_X | Araphid-pennate | 1589 |
| Stramenopiles | Ochrophyta | Chrysophyceae | Chrysophyceae_X | Chrysophyceae_Clade-E | 1615 |
| Stramenopiles | Ochrophyta | NA | NA | NA | 1697 |
| Alveolata | Ciliophora | Spirotrichea | Choreotrichida | NA | 2018 |
| Alveolata | Ciliophora | Litostomatea | Litostomatea_X | Litostomatea_XX | 2020 |
| Alveolata | Ciliophora | Spirotrichea | Choreotrichida | NA | 2033 |
| Alveolata | Dinoflagellata | Dinophyceae | Peridiniales | Thoracosphaeraceae | 2201 |
| Excavata | Discoba | NA | NA | NA | 2281 |

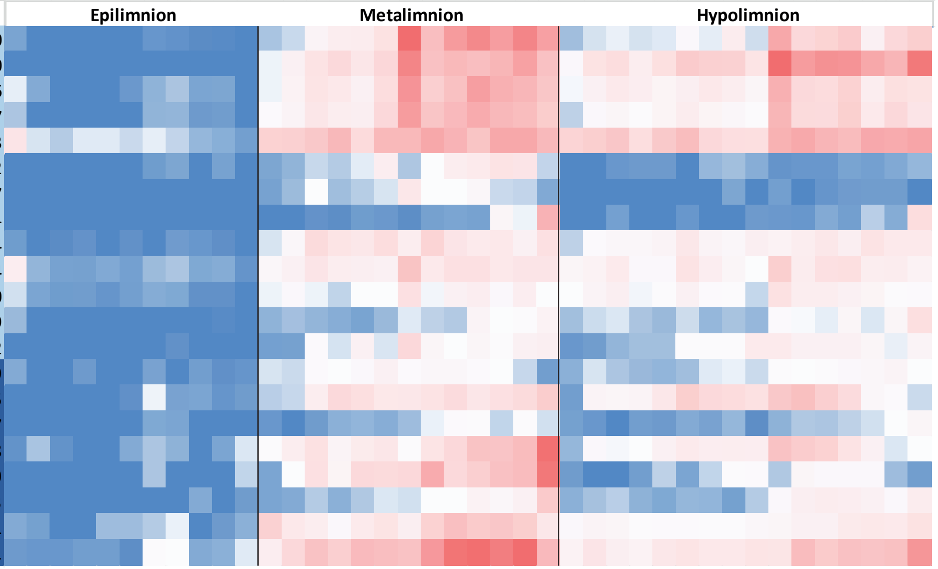


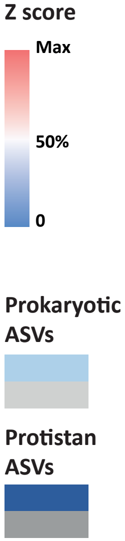


### Diverse small modules

| Phylum | Class | Order | Family | Genus | ID |
|---|---|---|---|---|---|
| Acidobacteriota | Vicinamibacteria | Vicinamibacterales | Vicinamibacteraceae | NA | 14 |
| Actinobacteriota | Acidimicrobiia | Microtrichales | Ilumatobacteraceae | CL500-29 marine group | 23 |
| Actinobacteriota | Acidimicrobiia | Microtrichales | Ilumatobacteraceae | CL500-29 marine group | 24 |
| Actinobacteriota | Actinobacteria | Frankiales | Sporichthyaceae | Candidatus Planktophila | 69 |
| Actinobacteriota | Actinobacteria | Frankiales | Sporichthyaceae | hgcI clade | 90 |
| Actinobacteriota | Thermoleophilia | Gaiellales | NA | NA | 165 |
| Bacteroidota | Bacteroidia | Flavobacteriales | Crocinitomicaceae | Fluviicola | 306 |
| Bacteroidota | Bacteroidia | Flavobacteriales | Crocinitomicaceae | Fluviicola | 307 |
| Bacteroidota | Bacteroidia | Flavobacteriales | Crocinitomicaceae | Fluviicola | 314 |
| Bacteroidota | Bacteroidia | Flavobacteriales | Flavobacteriaceae | Flavobacterium | 391 |
| Planctomycetota | Planctomycetes | Gemmatales | Gemmataceae | NA | 791 |
| Proteobacteria | Alphaproteobacteria | Sphingomonadales | Sphingomonadaceae | Sphingobium | 1087 |
| Proteobacteria | Gammaproteobacteria | Salinisphaerales | Solimonadaceae | NA | 1430 |
| Alveolata | Ciliophora | Litostomatea | Litostomatea_X | Litostomatea_XX | 2048 |
| Alveolata | Ciliophora | NA | NA | NA | 2039 |
| Stramenopiles | Ochrophyta | Chrysophyceae | Chrysophyceae_X | NA | 1614 |
| Stramenopiles | Ochrophyta | Chrysophyceae | Chrysophyceae_X | NA | 1629 |
| Rhizaria | Cercozoa | NA | NA | NA | 1900 |

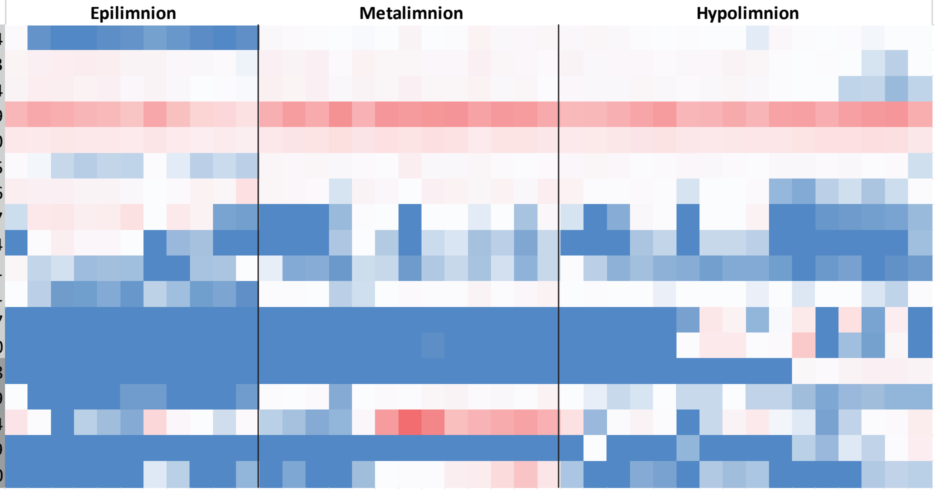
