## Supplementary figures and images for "Depth-dependent dynamics of protist communities as an integral part of spring succession in a freshwater reservoir"

### Additional File 9

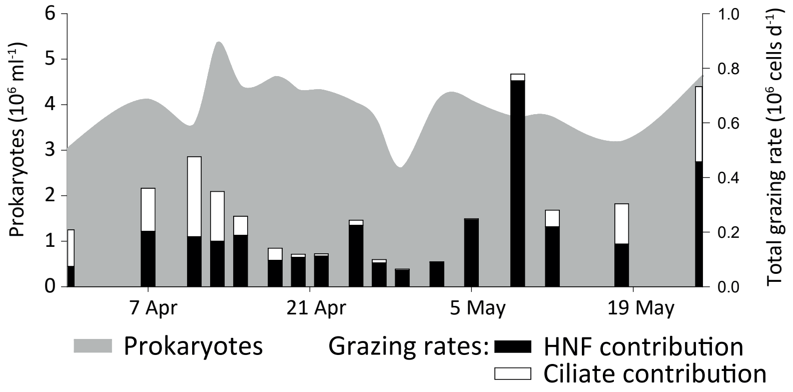


**Additional file 9:** Total grazing by protists in the epilimnion
